# Transcriptional memory is conferred by combined heritable maintenance and local removal of selective chromatin modifications

**DOI:** 10.1101/2023.12.16.571619

**Authors:** Pawel Mikulski, Sahar S.H. Tehrani, Anna Kogan, Izma Abdul-Zani, Emer Shell, Brent J. Ryan, Lars E.T. Jansen

## Abstract

Interferon-γ (IFNγ) transiently activates genes involved in inflammation and innate immunity. A subset of targets maintain a mitotically heritable memory of prior IFNγ exposure resulting in hyperactivation upon reexposure. Here we discovered that the active chromatin marks H3K4me1, H3K14Ac and H4K16Ac are established during IFNγ priming and selectively maintained on a cluster of GBP genes for at least 7 days in dividing cells in the absence of transcription. The histone acetyltransferase KAT7 is required for the accelerated GBP reactivation upon reexposure to IFNγ. In naïve cells, we find the GBP cluster is maintained in low-level repressive chromatin marked by H3K27me3 limiting priming in a PRC2-dependent manner. Unexpectedly, IFNγ results in transient accumulation of this repressive mark but is then selectively depleted from primed GBP genes during the memory phase facilitating hyperactivation of primed cells. Furthermore, we identified a cis-regulatory element that makes transient, long-range contacts across the GBP cluster and acts as a repressor, primarily to curb the hyperactivation of previously IFNγ-primed cells. Combined our results identify the putative chromatin basis for long-term transcriptional memory of interferon signalling that may contribute to enhanced innate immunity.

## Introduction

Cells exist in a dynamic environment, where they respond to a multitude of stimuli by rewiring their gene expression programmes. While acute transcriptional activation by external signals is well understood, the longer-term cellular consequences of such signals are poorly defined. Cells can maintain a memory of past stimulation that can be inherited for multiple cell generations. Post-stimulus epigenetic memory has been characterized mostly in the context of long-term gene repression. Prominent examples include read-write mechanisms that maintain gene silencing through DNA methylation, repressive histone modifications and Polycomb complex binding^1^. However, transient gene activation can also be memorized, known as long-term transcriptional memory^2^. Such memory of gene activation is relevant as it may alter the cellular response to future reexposure to activating signals.

Despite the existence of transcriptional memory phenomena across multiple cellular processes and species^3^, its molecular principles are obscure and represent an important gap in our understanding of gene expression regulation. Cellular exposure to the cytokine Interferon-gamma (IFNγ) is known to induce transcriptional memory and is an ideal model for discovering the underlying mechanisms. IFNγ induces a broad set of genes acting in inflammation, cell death, and host defence to pathogens and cancer^4^. In addition to the transient activation of a large number of genes, IFNγ induces long-term transcriptional memory of a subset of genes in different cell types, including innate immune macrophages, non-immune fibroblasts and cancer cells^5–8^. We and others discovered that genes that display memory tend to reside in genomic clusters^8^. One of these is a clustered family of genes encoding Guanylate-Binding Proteins (GBPs), GTPases that are crucial for inflammasome activation and protection against infections and cancer^9^. While IFNγ results in only a transient activation of GBPs, cells maintain a heritable epigenetic memory of activation for up to 14 days of continued proliferation in the absence of target gene expression^8^. This primed state results in hyperactivation of GBP genes upon re-exposure to IFNγ which may represent a crucial means for enhanced innate immune responses to repeated cellular insults.

To define the mechanism of transcriptional memory, we previously surveyed both trans-acting factors^8,10^ and chromatin features^8^. Here, we discovered an IFNγ-induced chromatin signature associated with transcriptional memory. This includes the acquisition of unique active histone marks (H3K4 monomethylation and H3K14 and H4K16 acetylation) and selective removal of repressive modification (H3K27 trimethylation). This chromatin signature is heritably maintained post-stimulation in proliferating cells, specifically at GBP genes that show transcriptional memory. After the initial IFNγ stimulation ceases and in the absence of ongoing transcription this heritable chromatin state functionally regulates future GBP gene expression triggered by IFNγ.

Unexpectedly, IFNγ-mediated induction of gene expression causes also a cluster-wide accumulation of repressive H3K27 trimethylation at and around GBP genes during initial stimulation. Subsequently, H3K27 trimethylation is removed selectively at GBP memory genes in primed and reinduction conditions. Furthermore, we uncovered a cis-regulatory element that generates long-range interactions within the GBP cluster enhanced during IFNγ induction. While these long-range interactions are not inherited post-stimulation, the element functions as a repressor of hyperactivation of GBP memory genes.

Our results are consistent with a model where the GBP memory loci selectively retain an epigenetically inherited chromatin signature after initial IFNγ stimulation, which in turn accelerates future expression hyperactivation upon IFNγ re-exposure. GBP gene hyperactivation is restricted by a repressive cis-regulatory element forming long-range interactions that are themselves not epigenetically inherited post-stimulation. Our results have broad mechanistic implications for the understanding of epigenetic memory of gene expression triggered by past exposure to the stimuli.

## Results

### Specific active chromatin modifications are established during priming and heritably maintained at GBP memory genes post-stimulation

To discover potential carriers of mitotically heritable memory of gene activation, we explored a previously established model of interferon-γ (IFNγ) gene activation^7,8^. We exposed human (HeLa) cells to recurrent IFNγ stimulation (priming and reinduction) separated by days or weeks in the absence of a stimulus and continuous cell proliferation (Fig. 1A). Analysis of our previously reported RNA-seq dataset^8^ revealed that the expression of the majority of IFNγ inducible genes are activated to a similar level, irrespective of whether they have been activated before, indicating no memory of prior exposure. However, a subset of genes shows hyperactivation of expression upon reinduction compared to priming (Fig. 1A, B), as described^5,8^. The most prominent among these is the GBP family of genes, where GBP1, GBP5 and GBP4 exhibit the highest degree of hyperactivation (strong memory GBPs), while GBP2 show weak hyperactivation. Interestingly, all these GBP genes are paralogs that are proximally arranged as a gene cluster on human chromosome 1. These findings suggest that an initial IFNγ stimulation (priming) induces a primed state within this cluster that allows faster and stronger expression of GBP memory genes upon IFNγ re-exposure (reinduction) (Fig. 1A, B). As the genes are not expressed in the intervening period between IFNγ pulses while cells continuously proliferate, this suggests that cells maintain an epigenetic mode of transcriptional memory.

**Fig. 1.**
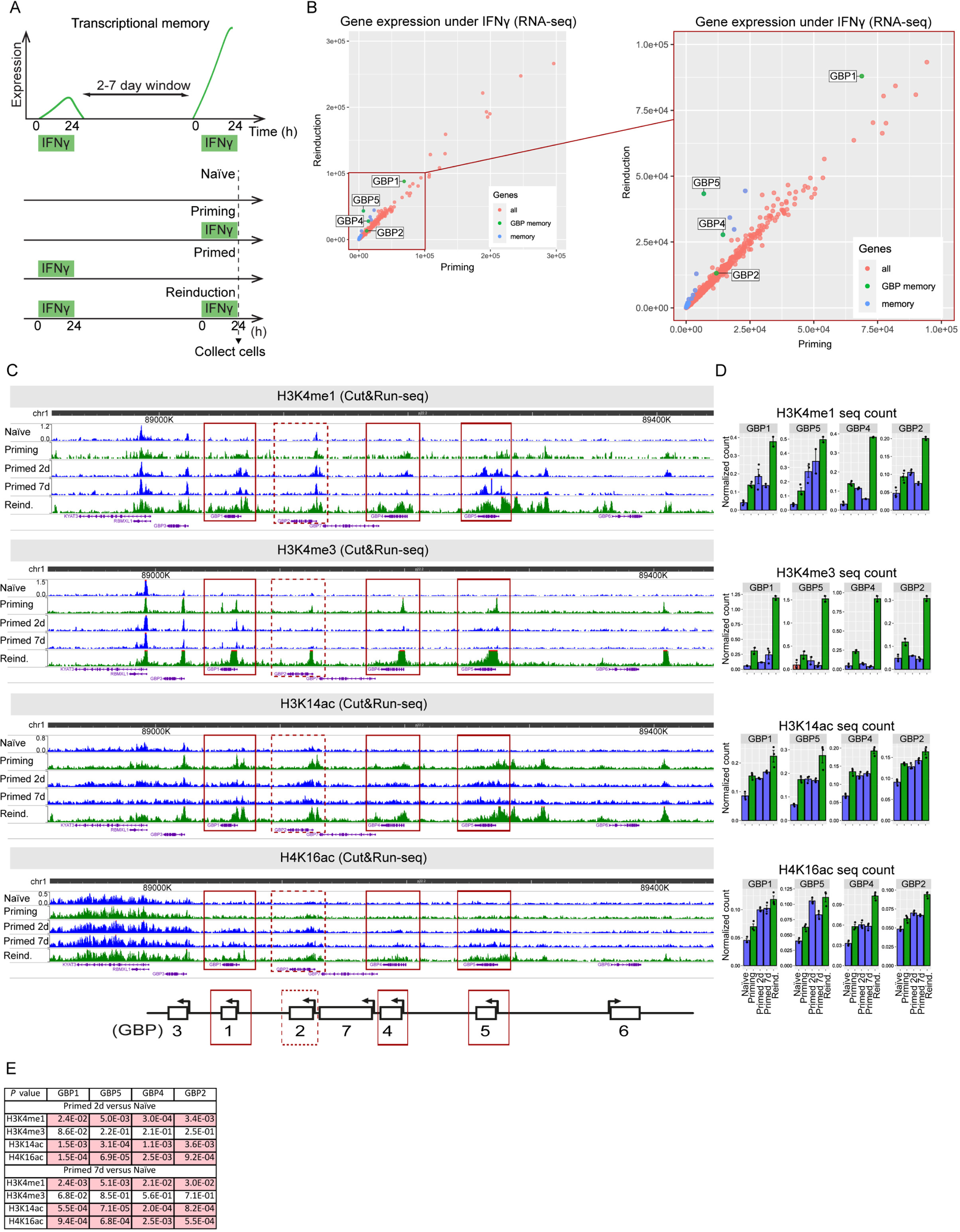
Specific active chromatin modifications are established during priming and heritably maintained at GBP memory genes post-stimulation. **A.** Experimental transcriptional memory regime outlining timing of IFNγ-incubation and cell harvesting. **B.** Gene expression plots for IFNγ-mediated stimulation comparing priming vs reinduction. Plot on the right represents detailed view from boxed area in left panel. Each dot corresponds to an individual IFNγ-stimulated gene, color-coded according to the legend. Data reanalyzed from^8^ **C.** Cut&Run-seq enrichment of active chromatin modifications in IFNγ-induced transcriptional memory regime represented as genome browser snapshots over GBP cluster. Red boxes indicate regions over GBP memory genes used for quantification. **D.** Quantification of normalized Cut&Run sequencing reads for respective chromatin modifications over GBP memory genes. The error bars correspond to SEM. The black dots on bar plots correspond to individual biological replicates. **E.** *P* values for relevant pairwise comparisons of quantifications are shown in panel D. *P* values ≤0.05 are highlighted in red. Statistical significance was calculated with two-sided *t*-test and prior determination of homo- or heteroscedasticity with F-test.

We sought to discover the mechanism driving transcriptional memory at GBP genes. We previously found that trans-acting factors including upstream IFNγ signalling, JAK kinase activity, Polymerase II occupancy and activation of the key downstream transcription factor STAT1 are not the carriers of long-term memory^10^. Similarly, cis-acting chromatin features that are associated with active gene expression, including an increase in chromatin accessibility, H3K27 acetylation and H3K36 trimethylation are only transiently associated with GBP genes but return to baseline levels upon loss of expression^8^.

Given the heritable nature of gene priming, we explored changes in specific chromatin modifications based on their potential to be maintained through cell division and their association with active genes that heritably control cell fate. These include histone H3K4 mono and trimethylation and H3K14 and H4K16 acetylation as they have been canonically implicated in regulating active chromatin states^11–14^. We assessed their enrichment in chromatin by Cut&Run-seq^15^ in naïve cells, during IFNγ stimulation, in the period post-stimulation (primed) and upon reinduction. While these modifications are present at basal levels in naïve cells, they accumulated at GBP promoters and bodies upon IFNγ induction and accumulated further upon reinduction (Fig. 1C, D), correlating with gene expression (Fig. 1A). The enhanced accumulation of these modifications relative to the level observed during priming at GBP genes is among the highest observed across the genome (Fig. S1A).

Upon IFNγ removal, H3K4me3 levels are rapidly lost in primed cells, with little remaining after 2 days and reach pre-stimulation levels by 7 days post IFNγ washout (Fig. 1C, D). This indicates H3K4me3 is an acute, non-memorized mark associated with ongoing expression that is reset when transcription ceases.

In striking contrast, we observed different dynamics of H3K4me1, H3K14ac and H4K16ac that were all maintained in primed cells at levels above those in naïve cells (Fig. 1C, D). The degree of retention is the highest on the strongest (GBP1, 4, 5) relative to weak GBP memory genes (GBP2) (Fig. 1C-E) or the rest of the genome (Fig. S1A). This suggests that H3K4me1, H3K14ac and H4K16ac are selectively retained on the chromatin of memory GBPs in primed cells, despite that lack of GBP expression and continuous cell proliferation for up to 7 days (∼7 cell divisions). These findings suggest that the maintenance of unique active chromatin modifications post-stimulation could confer transcriptional memory and allow differential expression of memory genes to recurrent stimulations.

### IFNγ-activated GBP cluster accumulates repressive chromatin that is selectively removed from GBP memory genes post-stimulation

In addition to the propagation of active chromatin marks, the removal of repressive marks may also contribute to the maintenance of a primed state. To investigate this, we analysed the Polycomb-mediated repressive histone mark, H3K27me3^16^, by Cut&Run-seq to assess chromatin enrichment during and after IFNγ stimulation. We find H3K27me3 is pre-established broadly across the GBP cluster in naïve cells, including intergenic regions, in agreement with the lack of GBP expression before stimulation (Fig. 2A-C, G; Fig. S2A-C). Surprisingly, while H3K27me3 has a known repressive role in transcription^16^, we find it further accumulates across the cluster during priming (Fig. 2A-C; Fig. S2), when GBP genes are upregulated. This broad accumulation is generally maintained over the cluster in primed and reinduction conditions (Fig. S2A-C). In contrast, H3K27me3 is locally depleted from gene bodies and proximal promoters of strong memory genes (GBP1, 2, 4, 5) (Fig. 2A-C). At these genes, loss of H3K27me3 is initiated after IFNγ priming and is maintained during memory and further depleted during reinduction. Genome-wide analysis reveals that the selective loss of H3K27me3 in primed and reinduction conditions is the most prominent at GBP memory genes (Fig. S2C).

**Fig. 2.**
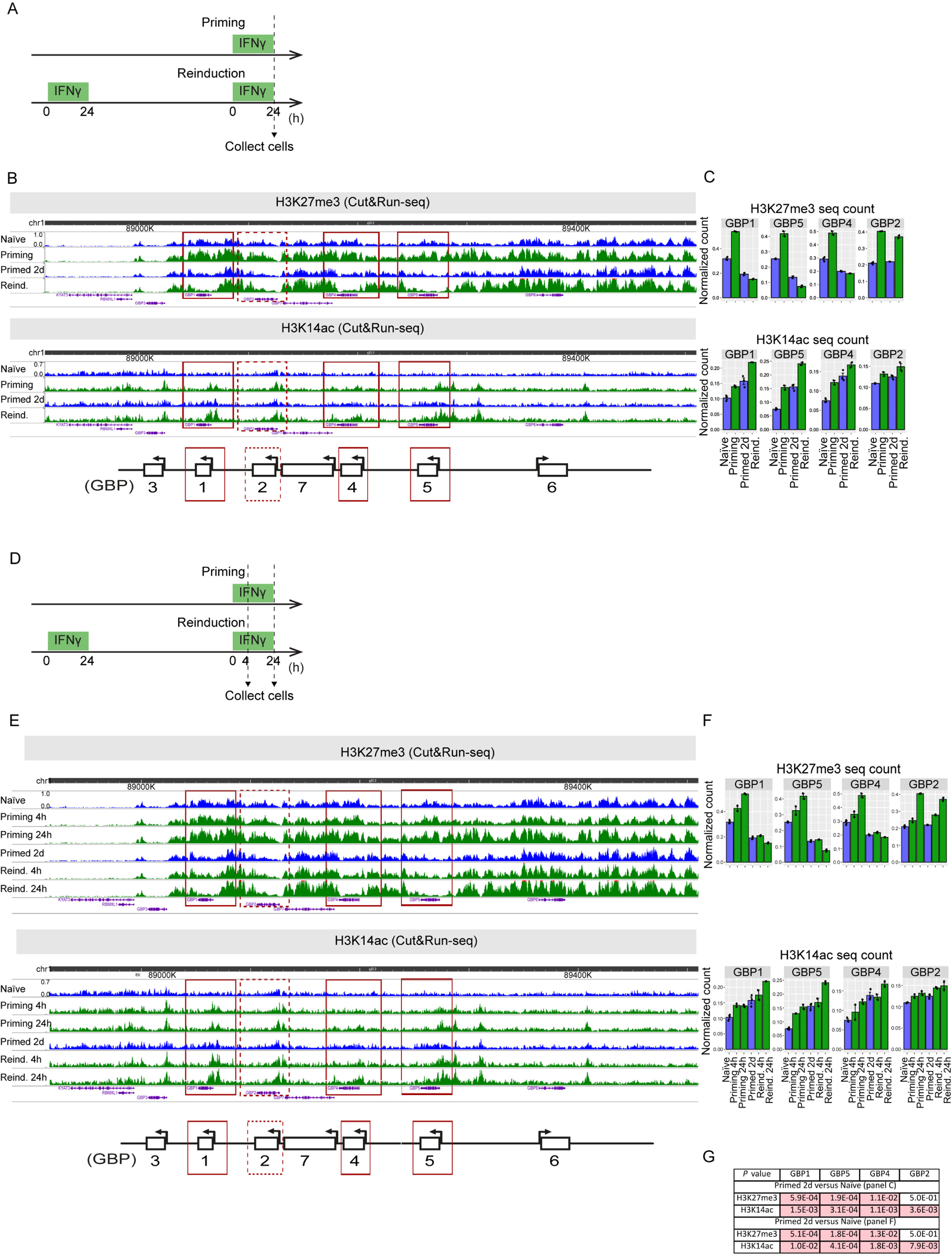
IFNγ-activated GBP cluster accumulates repressive chromatin that is selectively removed from GBP memory genes post-stimulation. **A.** Experimental transcriptional memory regime outlining timing of IFNγ-incubation and cell harvesting**. B.** Cut&Run-seq enrichment of indicated chromatin modifications during IFNγ-induced transcriptional memory regime represented as genome browser snapshots over GBP cluster. Red boxes indicate regions over GBP memory genes used for quantification. **C.** Quantification of normalized Cut&Run sequencing reads for respective chromatin modifications over GBP memory genes. **D.** Experimental transcriptional memory regime outlining timing of IFNγ-incubation and cell harvesting**. E.** Cut&Run-seq enrichment of indicated chromatin modifications during transcriptional memory regime with short (4h) and long (24h) IFNγ stimulation. The enrichment is represented as genome browser snapshots over GBP cluster. Red boxes indicate regions over GBP memory genes used for quantification. **F.** Quantification of normalized Cut&Run sequencing reads for respective chromatin modifications over GBP memory genes with short (4h) and long (24h) IFNγ stimulation. **G.** *P* values for relevant pairwise comparisons of quantifications shown in panels: C and F. Statistical significance was calculated with two-sided *t*-test and prior determination of homo- or heteroscedasticity with F-test. *P* values ≤0.05 are highlighted in red. The error bars on all bar plots in the figure correspond to SEM. The black dots on bar plots correspond to individual biological replicates.

Furthermore, we observed that the limits of this repressive domain coincide with previously reported TAD borders around the GBP cluster^17,18^ (Fig. S3A). H3K27me3 enrichment at GBP cluster overlaps with a B compartment as defined by HiC^17^ and shows overall higher levels compared to the cluster-wide active modifications, H3K14ac and H4K16ac (Fig. S3A). Such observation suggests that GBP cluster resides in a generally repressive chromatin domain.

To determine how repressive chromatin relates to active chromatin features at the GBP cluster we compared H3K27me3 with the H3K14ac mark within the same experiment as the latter is efficiently maintained at IFNγ-primed cells (Fig. 1C, D). Analysis 4 hours post IFNγ stimulation shows that both H3K14ac and H3K27me3 are already accumulated above naïve levels indicating that chromatin reorganization is rapid (Fig. 2D-G). Over the course of IFNγ activation, we observed a gradual increase in H3K14ac and a gradual local loss of H3K27me3 at GBP memory genes (Fig. 2D-G). The quantitative changes in chromatin structure may result either from a cell-autonomous gradual increase of target gene expression and/or an increase in the fraction of IFNγ-responsive cells^8^. Interestingly, we find that while during priming both H3K27me3 and H3K14ac accumulate on memory GBP genes, in primed cells, their local occupancy becomes antagonistic, where the highest H3K14ac peaks overlap with regions of local H3K27me3 depletion (Fig. 2A-D). This inverse enrichment is further extended during reinduction. These results suggest that while initially co-enriched, active and repressive chromatin modifications at the GBP cluster occupy locally distinct chromatin regions during the memory phase. The combined maintenance of active chromatin and selective removal of repressive modifications could underpin the transcriptional memory of GBP genes.e

### The writers for H3K14ac & H3K27me3 are functionally required for GBP gene expression and memory

To assess the functional requirement for the active retention of active chromatin and local loss of repressive histone marks in transcriptional memory at GBP genes, we depleted the writers for these marks in the context of IFNγ priming and memory. First, we targeted the H3K14ac writer, KAT7 (also known as MYST2/HBO1)^19,20^ as this mark is strongly maintained in the primed state, post-stimulation (Fig. 1C, D). To minimize off-target and indirect effects we transiently depleted KAT7 with two distinct siRNAs (KAT7 siRNA-1 and -2) during the memory phase, directly comparing naïve and primed cells (Fig. 3A). We assessed the expression level of GBP memory genes (GBP1, 4, 5) and non-memory controls (STAT1, IRF1) and KAT7 by a Real-Time quantitative PCR (RT-qPCR) (Fig. 3B. C).

**Fig. 3.**
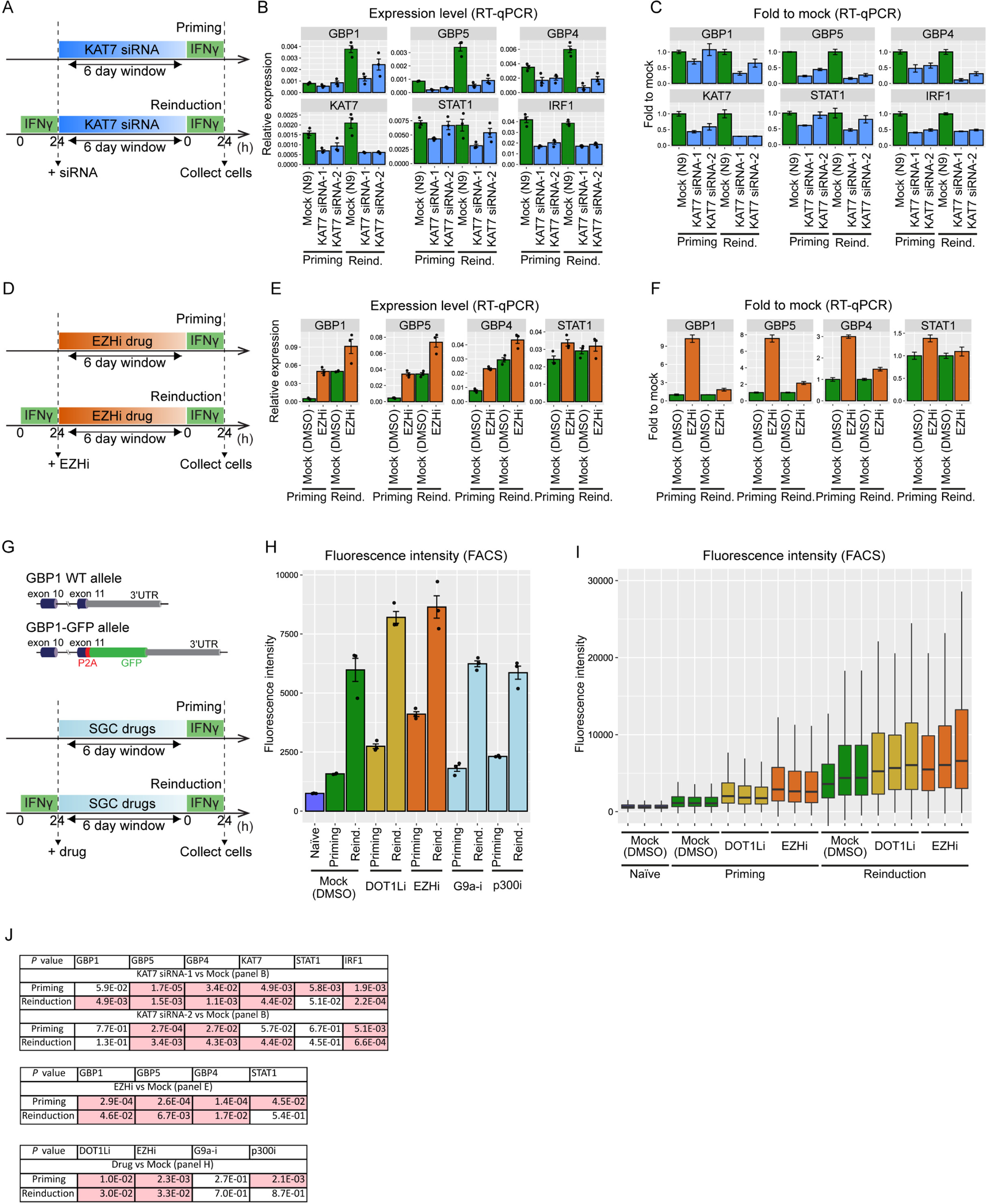
The writers for H3K14ac, H3K27me3 and H3K79me are functionally required for GBP gene expression and memory. **A.** Experimental scheme for transient KAT7 depletion during IFNγ priming or reinduction. **B.** Normalized RT-qPCR data of target genes upon IFNγ priming or reinduction after KAT7 depletion using two independent KAT7 siRNAs (siRNA-1 or -2) compared to mock siRNA control (N9). **C.** Fold changes between KAT7 siRNA-1 or -2 and mock control derived from RT-qPCR data in panel B. **D.** Experimental scheme for EZH1/2 inhibition during IFNγ priming or reinduction. **E** Normalized RT-qPCR data of target genes upon IFNγ priming or reinduction after EZH1/2 inhibition compared to mock control (DMSO). **F.** Fold changes between EZHi and mock control derived from RT-qPCR data in panel E. **G.** Top: Schematic outline endogenous GBP1 gene structure in GBP1-GFP reporter line. GFP reports GBP1 activation but is separated from GBP1 by a P2A ribosomal skipping peptide. Bottom: Scheme for secondary validation of putative regulators of IFNγ transcriptional memory from SGC small molecule screening in Fig. S4. **H.** Mean GBP1-GFP fluorescence intensities upon inhibition of putative regulators measured by FACS. Fluorescnece is assesed in mock control (DMSO), naïve, priming and reinduction conditions. **I.** Experiment shown in H, plotted as boxplots for DOT1Li and EZHi to show individual replicates and signal distribution across the cell population (minimum, 1^st^ quartile, median, 3rd quartile, maximum). **G.** *P* values for relevant pairwise comparisons of quantifications shown in panels: B, E and H. *P* values ≤0.05 are highlighted in red. Statistical significance was calculated with two-sided *t*-test and prior determination of homo- or heteroscedasticity with F-test. The error bars on all bar plots in the figure correspond to SEM. The black dots on bar plots correspond to individual biological replicates.

As expected KAT7 depletion generally results in downregulation of all tested genes (Fig. 3B, C). However, while the loss of expression of GBP memory genes was modest during priming, KAT7 depletion resulted in a stronger downregulation during reinduction (Fig. 3B, C, J). The stronger requirement for KAT7 during reinduction is specific for memory GBP genes as non-memory IFNγ-inducible controls (STAT1 and IRF1), showed a similar degree of downregulation in either condition (Fig. 3B, C). While we cannot exclude indirect effects, these results suggest that KAT7 and its catalytic product H3K14ac are functionally required to promote GBP memory gene expression, particularly during reinduction. This conditional dependency is consistent with the post-stimulus H3K14ac retention and high enrichment in primed and reinduction states (Fig. 1C, D; Fig. 2A-C).

Next, we targeted EZH1/2, the methyltransferases of the PRC2 complex that generate the repressive H3K27me3 mark^21,22^. To target this complex with high temporal control we took advantage of a selective small molecule inhibitor (EZHi – UNC1999)^23^ in an experimental regime similar to that of KAT7 inhibition (Fig. 3D). EZH1/2 inhibition resulted in the upregulation of all tested genes (Fig. 3E, F), consistent with its known role in gene repression. Interestingly, the upregulation of GBP memory genes is much stronger during priming than reinduction (Fig. 3E, F; Fig. S4A, B). In contrast, non-memory IFNγ target genes showed no significant upregulation compared to mock (STAT1) in either condition (Fig. 3E, F; Fig. S4A, B). These results indicate that EZH1/2, likely through its product, H3K27me3, represses the expression of GBP memory genes, particularly during priming. This conditional dependency is consistent with the high enrichment of H3K27me3 in the GBP cluster in naïve cells and during priming. In primed cells and upon reinduction, H3K27me3 is largely depleted, consistent with a minor functional role for EZH1/2 in expression at this stage (Fig. 2A-C).

Combined our manipulation of KAT7 and EZH1/2 suggests that both the active and repressive chromatin dynamics we observed during priming and memory are functionally required for GBP expression and memory of the primed state.

### Small molecule screening identifies putative regulators of IFNγ-induced transcriptional memory

To discover any other potential regulators of IFNγ priming of GBP genes, we screened a subset of the Epigenetic Chemical Probe Library from the Structural Genomics Consortium’s (SGC)^24^ containing 21 small molecules targeting chromatin modifiers. To assess these compounds, we built an IFNγ expression and memory reporter cell line in which we included GFP as part of the endogenous GBP1 mRNA, expressed as a separate polypeptide, fatefully reporting GBP1 expression in HeLa cells (Fig. S4C). Next, we treated these cells with the indicated SGC compounds before IFNγ activation of the GBP1 reporter and screened for GFP fluorescence by high throughput microscopy (Fig. S4C). While most compounds did not significantly affect IFNγ induction of GBP1-GFP we identified SGC0946 (inhibitor for DOT1L – H3K79 methyltransferase)^25^ as a putative hit that leads to enhanced GBP1-GFP expression during IFNγ activation (Fig. S4D). We also observed a small, but significant, GBP1-GFP downregulation with NVS-MLLT-1 ((inhibitor for MLLT1 – chromatin reader component of super elongation complex)^26^ and SGC6870 (inhibitor for PRMT6 – arginine methyltransferase)^27^. We also identified UNC1999, the inhibitor for EZH1/2 in this screen validating our earlier RT-qPCR results on EZHi (Fig. S4D, E; Fig. 3D-F).

We then explored the potential role of the positive hits (DOT1Li and EZHi) in priming and reinduction. Furthermore, to assess whether the effect of KAT7 depletion on GBP expression shown above is specific, we also included an inhibitor for p300 (p300i), an acetyltransferase for H3 and H4 distinct of KAT7^28^, and an inhibitor for G9a (G9a-i), a methyltransferase for repressive H3K9 methylation^29^. We measured GBP1-GFP expression during priming and reinduction following drug treatments by FACS which allowed us to score a large number of cells (Fig. 3G). We found that treatments with DOT1Li and EZHi increased GBP1-GFP fluorescence in both conditions (Fig. 3H, I). In contrast, inhibition of G9a did not alter GBP1-GFP expression, suggesting that GBP expression is selectively dependent on Polycomb-mediated repressive chromatin and not H3K9 methylation or its writer (Fig. 3H). Similarly, inhibition of p300 led only to a small, but significant, change in GBP expression, suggesting that the histone acetylation installed by KAT7 is selectively required for GBP expression rather than p300-mediated acetylation such as H3K27Ac (Fig. 3H). Importantly, in agreement with RT-qPCR results (Fig. 3D-F), we observed a higher degree of GBP1-GFP upregulation in priming than reinduction (Fig. 3H, I) upon EZH1/2 inhibition, confirming that the H3K27me3 and/or EZH1/2 are key limiting factors, particularly during GBP priming.

In sum, by small molecule screening, we identified EZH1/2, but also DOT1L, as a potential repressor of GBP expression memory genes during IFNγ stimulation. These findings suggest that, in addition to the repressive role of H3K27me3 described above, DOT1L or its catalytic products H3K79me1/2/3 may contribute to GBP cluster control, although this requires future investigation.

### A cis-regulatory element controls gene repression across the GBP cluster

Further analysis of the distribution of active and repressive chromatin modifications revealed their accumulation not only at GBP genes themselves but also at intergenic regions across the cluster (Fig. 1; Fig. 2). This suggests that the GBP genes and the surrounding chromatin domain may be regulated globally across the cluster by a common control element.

To discover such elements, we compared the enrichment of H3K4me1, H3K14ac, H4K16ac with the binding of the key IFNγ-induced transcription factor STAT1^4^. In addition to GBP gene promoters, we previously found STAT1 to target two elements 16 and 37 kb upstream of the GBP5 promoter (Fig. 4A)^10^. Strikingly, we find high enrichment of active chromatin modifications (H3K4me1, H3K14ac, H4K16ac) in primed cells not only at the bodies and promoters of GBP genes but also at the STAT1-bound intergenic elements, termed Element 1 (E1) and Element 2 (E2) (Fig. 4B, D). Similar to GBP gene promoters, these elements also exhibit removal of H3K27me3 in primed cells (Fig. 4C, D) and generally display similar chromatin landscape dynamics throughout IFNγ priming regime as GBP memory genes (Fig. 1; Fig. 2; Fig. 4B, C).

**Fig. 4.**
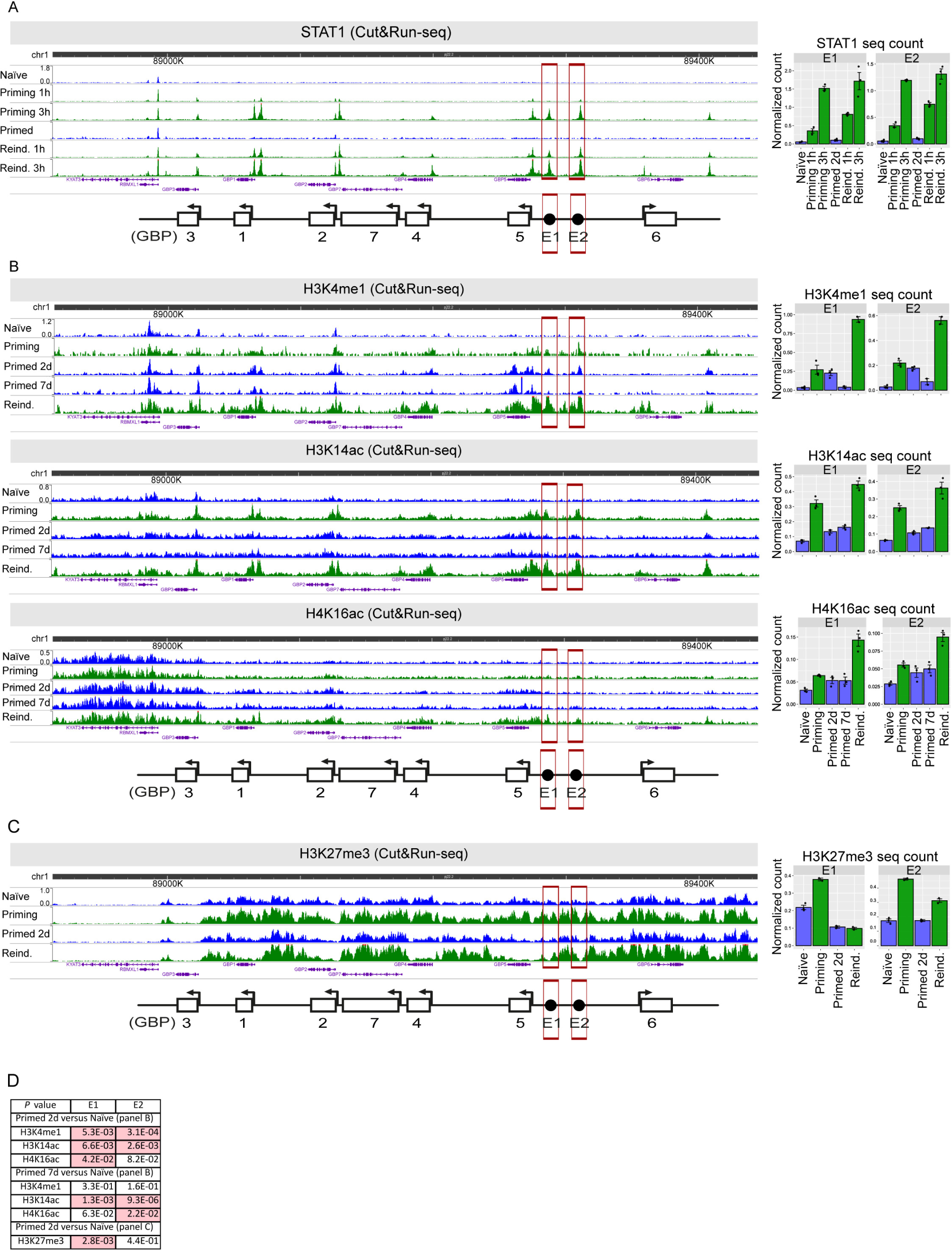
GBP cluster contains uncharacterized, transcription factor-bound cis-regulatory elements with a transcriptional memory chromatin signature. **A.** Cut&Run-seq of the IFNγ-activated transcription factor STAT1 after 0, 1h and 3h of activation in niave and primed cells. Genome browser snapshots over GBP cluster (left panel) and quantification of normalized Cut&Run sequencing reads over identified cis-regulatory elements (right panel). Data reanalzyed from^10^. **B.** Cut&Run-seq enrichment of active chromatin modifications during IFNγ-induced transcriptional memory regime (as shown in Figure 1) represented as genome browser snapshots over GBP cluster (left panel) and quantification of normalized Cut&Run sequencing reads over identified cis-regulatory elements (right panel). **C.** As in B but for the repressive chromatin modification H3K27me3. Red frames on all panels indicate regions used for quantification. **D.** *P* values for relevant pairwise comparisons of quantifications shown in panels: A, B and C. *P* values ≤0.05 are highlighted in red. Statistical significance was calculated with two-sided *t*-test and prior determination of homo- or heteroscedasticity with F-test. The error bars on all bar plots in the figure correspond to SEM. The black dots on bar plots correspond to individual biological replicates.

To determine the functional relevance of these putative cis-regulatory elements, we generated CRISPR knock-out lines for E1 and E2 and analyzed the consequences for IFNγ priming and reinduction of GBP genes by RT-qPCR (Fig. 5A). First, we assessed a polyclonal knockout line for E1 and two independently generated polyclonal knockout lines for E2 (*E2-1, E2-2,* Fig. 5A). While loss of E1 does not have a significant effect on GBP expression, E2-1 and E2-2 showed a marked upregulation of GBP memory genes, primarily upon reinduction (Fig. 5B). To confirm these results and exclude clonal heterogeneity, we subcloned a monoclonal line from the *E2-2* population and subjected it to a full IFNγ stimulation regime alongside wild type controls (Fig. 5C). In agreement with the polyclonal lines, we find that loss of E2 results in strong upregulation of memory GBP genes, selectively during IFNγ reinduction while its contribution to initial priming is modest (Fig. 5D). This effect is specific to strong memory GBP genes (GBP1, 4 and 5) as STAT1 is not affected and weak memory control GBP2 is only marginally affected (Fig. 5C, D).

**Fig. 5.**
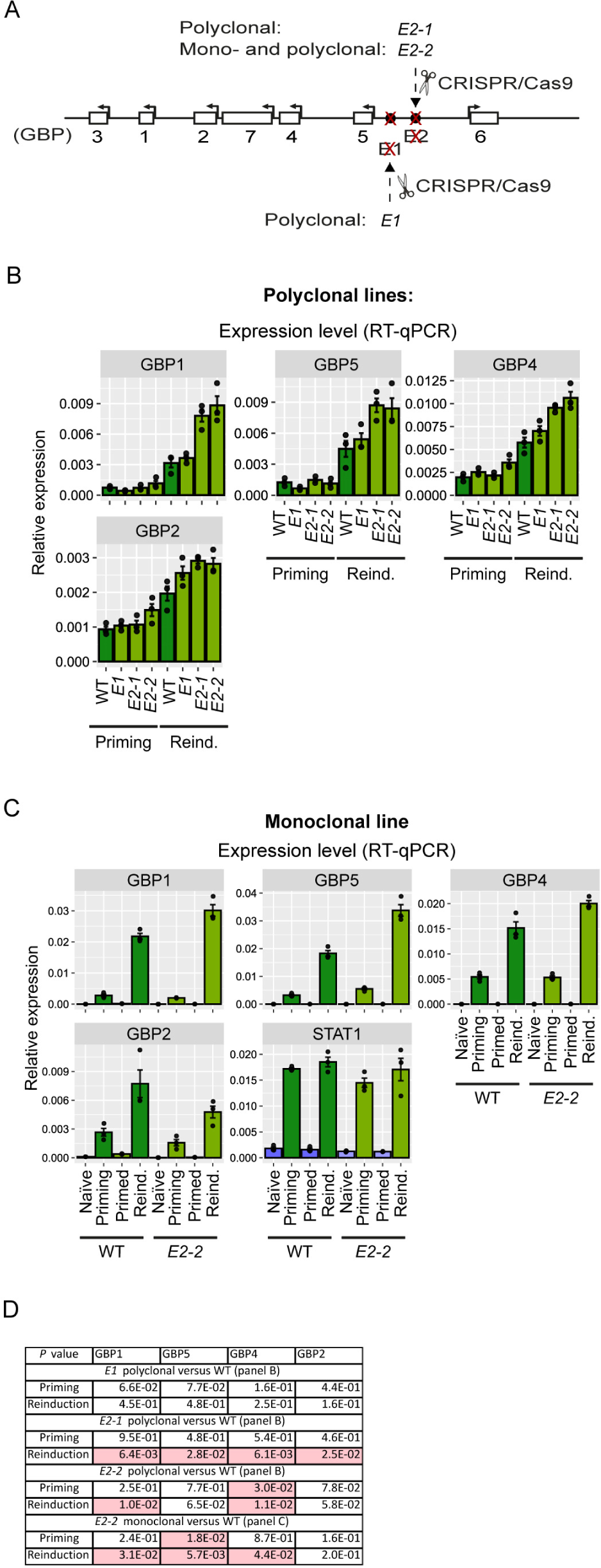
The E2 cis-regulatory element controls gene repression across the GBP cluster. **A.** Schematic of GBP gene cluster with indicated CRISPR deletions of E1 and E2, either as polyclonal or monoclonal population. E2-1 and E2-2 indicate two independent deletions generated with distinct gRNAs (see methods). Monoclonal line was isolated from E2-2 polyclonal line by FACS. **B.** Expression of GBP memory genes in wildtype (WT) and polyclonal E1, E2-1, E2-2 deletion lines during IFNγ priming or reinduction measured by RT-qPCR. **C.** Expression of GBP memory genes in wildtype (WT) and monoclonal E2-2 deletion line measured by RT-qPCR following IFNγ-stimulation regime: naïve, priming, primed and reinduction conditions (as Figure 1A). The error bars on all bar plots in the figure correspond to SEM. The black dots on bar plots correspond to individual biological replicates. **D.** *P* values for relevant pairwise comparisons of quantifications shown in panels: B and C. *P* values ≤0.05 are highlighted in red. Statistical significance was calculated with two-sided *t*-test and prior determination of homo- or heteroscedasticity with F-test. The error bars on all bar plots in the figure correspond to SEM. The black dots on bar plots correspond to individual biological replicates.

In sum, we discovered that the E2 cis-regulatory element is a transcriptional repressor of GBP genes, not only of the proximal GBP5 gene but across the GBP cluster including distant loci (i.e. GBP1). Our finding that a role for E2 is more pronounced in primed cells than in naïve conditions, suggests a selective role in transcriptional memory, curbing hyperexpression of GBPs.

### Cis-regulatory elements mediate cluster-wide interactions upon IFNγ stimulation

Our discovery of the cluster-wide role of E2 in repressing GBP expression suggests that it may act through long-range interactions. Chromatin looping e.g. in the context of enhancers or silencers contacting gene promoters has been previously implicated in epigenetic memory^30–33^. To explore this possibility, we employed Capture-C^34^, a modified HiC-type chromosome conformation capture method allowing unbiased assessment of all chromatin interactions from a selected genomic viewpoint. We designed specific hybridization probes to isolate the E2 locus following *in vivo* crosslinking, fragmentation and self-ligation to identify distal chromatin interactions. We also isolated a Cohesin-enriched locus upstream of GBP6 (labelled CH-C) that we previously identified as a repressor of the GBP memory genes^8^. We performed Capture-C in the context of the IFNγ stimulation regime and found that both E2 and CH-C loci broadly engage chromatin, selectively within the GBP cluster across all conditions (Fig. 6A). The boundaries of these interactions coincide with previously identified Cohesin-enriched sites^8^ and TAD borders in naïve Hela cells^17^. These interaction boundaries also overlap with the delimited enrichment of heritable chromatin modifications (Fig. 1; Fig. 2; Fig. S3), suggesting that the GBP cluster forms a specific chromatin domain distinct from neighbouring regions. Indeed, we did not detect interactions beyond the GBP locus on chromosome 1, nor trans-interactions to other chromosomes (Fig. 6A, S5A).

**Fig. 6.**
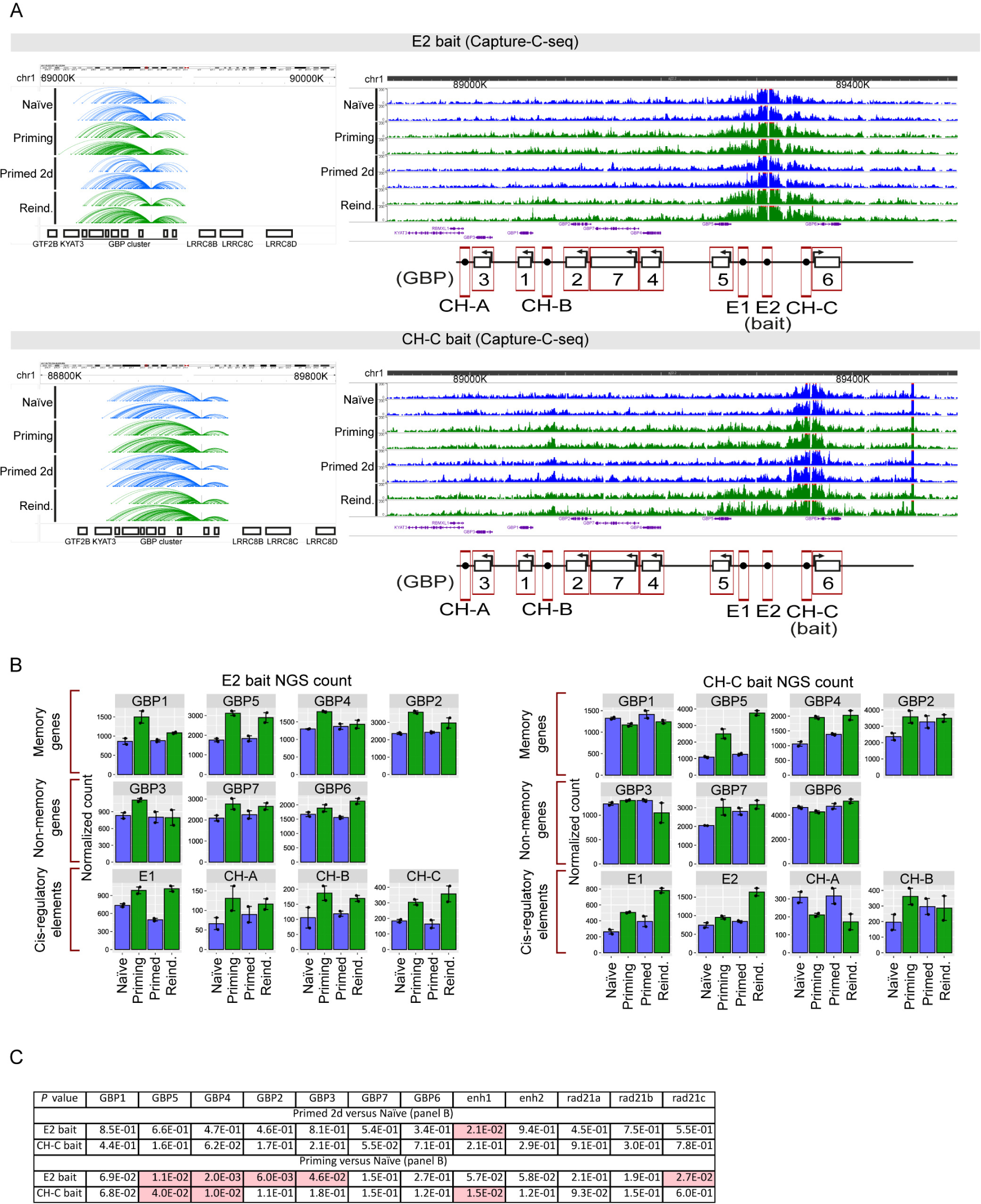
Cis-regulatory elements mediate cluster-wide interactions enhanced during IFNγ stimulation. **A.** Capture-C data showing long-range interactions from element E2 (top panel) or Cohesin site CH-C (bottom panel). The results show zoom-out (left panel) and zoom-in (right panel) genome browser snapshots from normalized Capture-C sequencing reads at and around GBP cluster. Genome browser tracks show 2 biological replicates per condition during IFNγ-stimulation regime. Red boxes correspond to the regions used for read quantification and the baits. **B.** Quantification of normalized Capture-C sequencing reads for E2 (left panel) or CH-C (right panel) baits across the GBP cluster: GBP memory genes (GBP1, 4, 5), GBP non-memory genes (GBP3, 6, 7, GBP1P1) and cis-regulatory elements (E1, E2, Cohesin sites (CH-A, CH-B, CH-C)). **C.** *P* values for relevant pairwise comparisons of quantifications shown in panel B. *P* values ≤0.05 are highlighted in red. Statistical significance was calculated with two-sided *t*-test and prior determination of homo- or heteroscedasticity with F-test. The error bars on all bar plots in the figure correspond to SEM. The black dots on bar plots correspond to individual biological replicates.

Further analysis of interactions within the GBP cluster revealed that, while interactions occur in the absence of GBP gene expression (naïve cells), IFNγ priming of the cluster triggers a marked increase in contact frequency (Fig. 6A, B), suggesting that IFNγ triggers spatial compaction of the cluster. Both the E2 and CH-C loci show enhanced engagement with virtually all genes and loci tested within the cluster. However, these contacts are transient and occur only during IFNγ activation and are reset in primed cells (Fig. 6). Consistent with this, we find no memory of long-range contact resulting in a similar degree of long-range engagement of E2 and CH-C with the GBP cluster upon reinduction (Fig. 6).

We validated these Capture-C results by conventional 3C-qPCR experiments. Using primers probing E2 or CH-C in combination with selected regions within the GBP cluster. We confirmed that pairwise interactions between these loci or with GBP promoters are increased upon IFNγ activation of the cluster but reset upon IFNγ withdrawal (Fig. S6A), while no ligation controls and distal loci confirm the selectivity of the method (Fig. S7).

Combined, we identified long-range interactions within the GBP cluster, delimited by Cohesin-marked boundaries. We find that the long-range interactions are associated with an acute non-memorized response to IFNγ. Importantly, in addition to contacting GBP genes, we also identified increased interactions between the E2 and CH loci (Fig. S6), suggesting that both repressive cis-regulatory elements could potentially regulate each other.

### Delayed activation of repressive cis-regulatory elements facilitates hyperexpression of GBP memory genes following IFNγ priming

The GBP cluster is rapidly activated by IFNγ and primed for hyperactivation upon reinduction (Fig. 1A, B). Yet, paradoxically, we identified a novel cis-regulatory element that represses GBP expression (Fig. 5). We next explored how the GBP cluster can be strongly activated despite the repressive effect of the E2 element. As E1 and E2 are STAT1 transcription factor-bound (Fig. 4A) we expected these elements to produce non-coding RNAs as commonly found at enhancer elements^35^. Indeed, we detected ncRNAs by RT-qPCR specifically upon IFNγ activation (Fig. S8) and used these as a readout of the activity of these elements. Importantly, both cis-regulatory elements show hyperactivation during IFNγ reinduction compared to priming (Fig. S8), indicating they exhibit transcriptional memory, similar to GBP memory genes.

We hypothesized that the GBP genes and the E1 and 2 elements are activated with different kinetics allowing GBP genes to be rapidly activated but their expression curbed at a later stage. To test this, we performed a timecourse experiment in which we assessed the expression of both E1 and E2 as well as GBP genes at an early (4h) and later (24h) timepoint (Fig. 7A). We confirmed that both E1, E2 and GBP genes show transcriptional memory with enhanced expression upon re-exposure to IFNγ (Fig. 7B). However, they show a striking difference in the dynamics of activation. The GBP memory genes, while poorly expressed during priming, show a very rapid activation upon reinduction (Fig. 7B). Already at 4 hours of IFNγ, primed cells show much higher expression than at any time during priming. In contrast, the E1 and E2 elements showed a marked delay in reactivation. At 4 hours both E1 and E2 are expressed at levels much lower than their priming levels (Fig. 7B). These results indicate that the cis-regulatory elements exhibit a delay of IFNγ-mediated transcriptional activation compared to GBP memory genes.

**Fig. 7.**
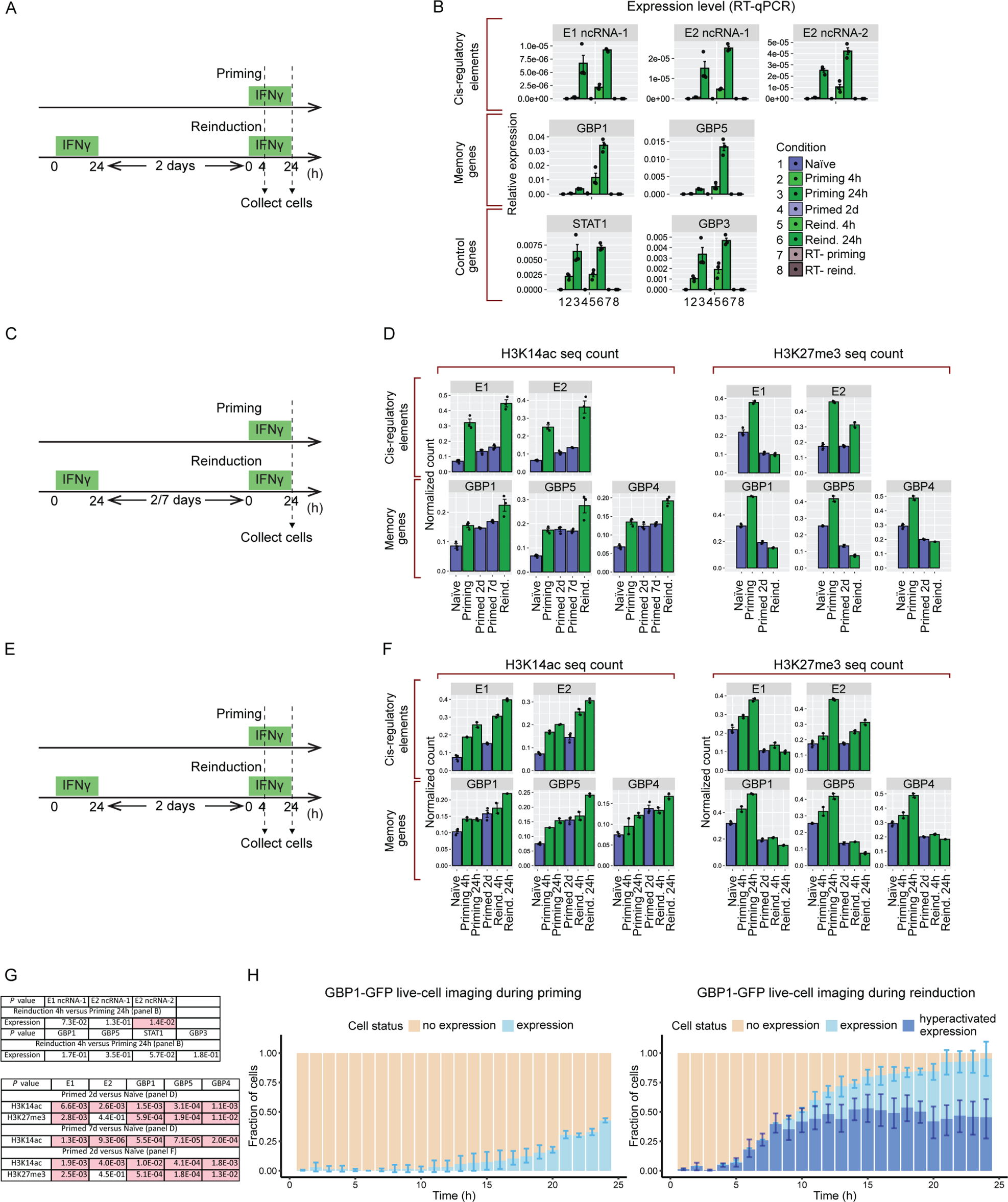
Delayed activation of cis-regulatory elements facilitates hyperactivation of GBP memory genes following IFNγ priming. **A.** Experimental transcriptional memory regime outlining timing of IFNγ-incubation and cell harvesting**. B.** Expression levels of target loci (cis-regulatory elements, GBP memory genes, control non-memory genes) measured by RT-qPCR following IFNγ-stimulation regime: naïve, priming, primed and reinduction conditions. cDNA synthesis negative controls [RT-(no reverse transcriptase)] are included for priming and reinduction conditions. **C.** Experimental transcriptional memory regime outlining timing of IFNγ-incubation and cell harvesting**. D.** Comparative quantification of H3K14ac (left panel) and H3K27me3 (right panel) enrichment between cis-regulatory elements and GBP memory genes. The results correspond to Cut&Run sequencing read count presented in Fig. 1, 2 and 4. **E.** Experimental transcriptional memory regime outlining timing of IFNγ-incubation and cell harvesting**. F.** Comparative quantification of H3K14ac (left panel) and H3K27me3 (right panel) enrichment between cis-regulatory elements and GBP memory genes in transcriptional memory time course. The results correspond to Cut&Run sequencing read count presented separately in Fig. 2C, D. **G.** *P* values for relevant pairwise comparisons of quantifications shown in panels: B, D and F. *P* values ≤0.05 are highlighted in red. Statistical significance was calculated with two-sided *t*-test and prior determination of homo- or heteroscedasticity with F-test. The error bars on all bar plots in the figure correspond to SEM. The black dots on bar plots correspond to individual biological replicates. **H.** Time-lapse of live-cell GBP1-GFP protein expression during priming (left) and reinduction (6 days after priming) (right). The fraction of cells with with expression and hyperactivated expression of GBP1-GFP is plotted for each time point. Hyperactivated expression during reinduction is defined as levels above those observed during priming (see methods). The bars represent mean of three replicates. The error bars correspond to SD.

Interestingly, when comparing the chromatin signatures of E1 and 2 with that of GBP genes we observed that the cis-regulatory elements exhibit a marked loss of H3K4me1, H3K14ac, H4K16ac relative to the levels established during priming whereas GBP genes maintain these marks to levels similar or even higher than those established during IFNγ priming (Fig. 7C, D), consistent with their more rapid reactivation upon IFNγ reinduction. H3K27me3 was lost from both E1 and GBP genes in primed cells and upon reinduction. However, we noticed that the E2 element does not show such a pronounced loss of H3K27me3 (Fig. 7C, D), suggesting that E2 maintains a more repressive chromatin signature.

Overall, the delayed expression of the E1 and 2 elements, weaker maintenance of active chromatin marks and the retention of repressive H3K27me3 at E2 may explain how GBP memory genes can initially ‘escape’ from its repressive function.

We hypothesise that at a later stage during IFNg re-exposure, the E2 element acts to curb GBP activation, preventing excessive hyperactivation. To test this, we turned to our GBP1-GFP reporter and monitored GFP expression during the 24-hour of priming and reinduction by IFNg by live-cell imaging (Fig. S8). Consistent with our earlier FACS readouts (Fig. 3), GBP1-GFP is activated to modest levels during priming but hyperactivated during reinduction of previously primed cells (Fig. 7H; S8D). We scored the number of cells that either activated GBP1-GFP to levels observed during priming or showed hyperactivation of GBP1-GFP unique to primed cells. We observed that the number of cells expressing GBP1 during priming gradually increases over the 24-hour period (Fig. 7H). In contrast, in primed cells the number of cells that hyperactivate GBP1, while initially increasing, plateaus after approximately 10 hours (Fig. 7H). This temporal dynamics is consistent with the expression dynamics and chromatin status of the cis-regulatory elements in the GBP cluster whose delayed activation allows initial rapid GBP induction but limits later hyperactivation.

## Discussion

Long-term transcriptional memory phenomena have been observed in species ranging from yeast to plants to humans^3^. In mammals, the heritable priming of cells by IFNγ is shown to last for several weeks through multiple cell division cycles, in the absence of ongoing transcription^5,7,8^. Despite its strong epigenetic nature, the molecular basis for what carries this memory has remained elusive.

We have now identified a set of chromatin modifications (H3K4me1, H3K14ac, H4K16ac) that are established on the GBP gene cluster which shows strong transcriptional memory. These marks are maintained for at least 7 days during which cells undergo multiple rounds of genome duplication and cell division. Importantly, these marks are associated with promoting transcription yet are maintained in the absence of detectable target gene expression making them putative carriers of memory. Their stable maintenance in proliferating cells in the absence of the initial trigger, suggests active propagation allowing the re-establishment of these chromatin modifications post-replication. Read-write mechanisms engaging in a feedback loop have been described for repressive marks such as Polycomb-mediated H3K27me3^36,37^, H3K9 methylation at heterochromatin^38^, as well as for DNA CpG methylation^39,40^.

Specific active chromatin modifications including those we identified here, are reported to be locally maintained on mitotic chromosomes, constituting ‘mitotic bookmarks’^41–43^. This behaviour is consistent with a role as mediators of transcriptional memory. However, how they engage in read-write feedback to avoid dilution during cell division remains unclear and is an important future direction of inquiry. Furthermore, our results are also consistent with the previously identified role of H3K4me1 in enhancer priming^44–46^ indicating this mark can be stably maintained.

In addition to the maintenance of active chromatin, we find that the PRC2 mark H3K27me3 is established across the GBP gene cluster during priming. It is unclear why strong transcriptional activation of the GBP cluster results in the recruitment of both active as well as repressive chromatin. This repsonse may be part of a mechanism to limit the otherwise strong and rapid activation of IFNγ targets. This is consistent with our finding that limiting PRC2 activity, that is responsible for H3K27 methylation, results in GBP hyperactivation. Importantly, H3K27me3 repressive chromatin is selectively depleted from memory genes and maintained at a low level in primed cells. The failure to re-establish H3K27me3 following IFNγ priming may be important to the priming of GBP genes. We propose that transcriptional memory is a consequence of the combined maintenance of active marks with the selective loss of repressive marks established during priming.

Furthermore, we identified a novel cis-regulatory element within the GBP cluster that makes extensive contacts with GBP genes across the cluster during IFNγ gene activation. These long-range contacts are not themselves inherited which indicates they are a consequence rather than the cause of transcriptional activation. Importantly, we discovered that the E2 element exerts a repressive effect on GBP expression, particularly during extended exposure to IFNγ reactivation. The E2 element has the signatures of an enhancer that is bound by the STAT1 transcription factor and generates RNAs during activation, yet, in the context of GBP expression, it acts as a repressive element. IFNγ target genes are proinflammatory and GBP gene expression has been shown to affect cell viability^47^. We postulate that the E2 element is important to prevent excessive GBP activity, particularly in cells already primed by IFNγ. We previously identified a Cohesin-bound boundary element of the GBP cluster that we found to have a similar repressive effect on GBP hyperactivation^8^. Possibly E2 and this boundary element cooperate in this function. How these elements exert their repressive effect on GBP expression remains an open question.

In summary, our findings suggest that transcriptional memory is mediated by a balance of unique active and repressive chromatin modifications that are differentially inherited in IFNg-primed cells, resulting in memory of prior IFNg exposure (Fig. 8). The output of transcriptional memory is regulated by cis-regulatory elements but their IFNg-dependent long-range contacts are not inherited as an epigenetic memory factors in IFNg-primed cells (Fig. 8).Interferons are important mediators of innate and adaptive immunity and are central to the priming of innate immune cells^48^. Key effectors of interferon signalling such as macrophages^48^ play a role in trained immunity where the organisms maintain a long-term memory of prior immune activation^49^. The mechanisms we uncover here may contribute to the cellular memory of innate immune signals that underpin the physiological state of trained immunity.

**Fig. 8.**
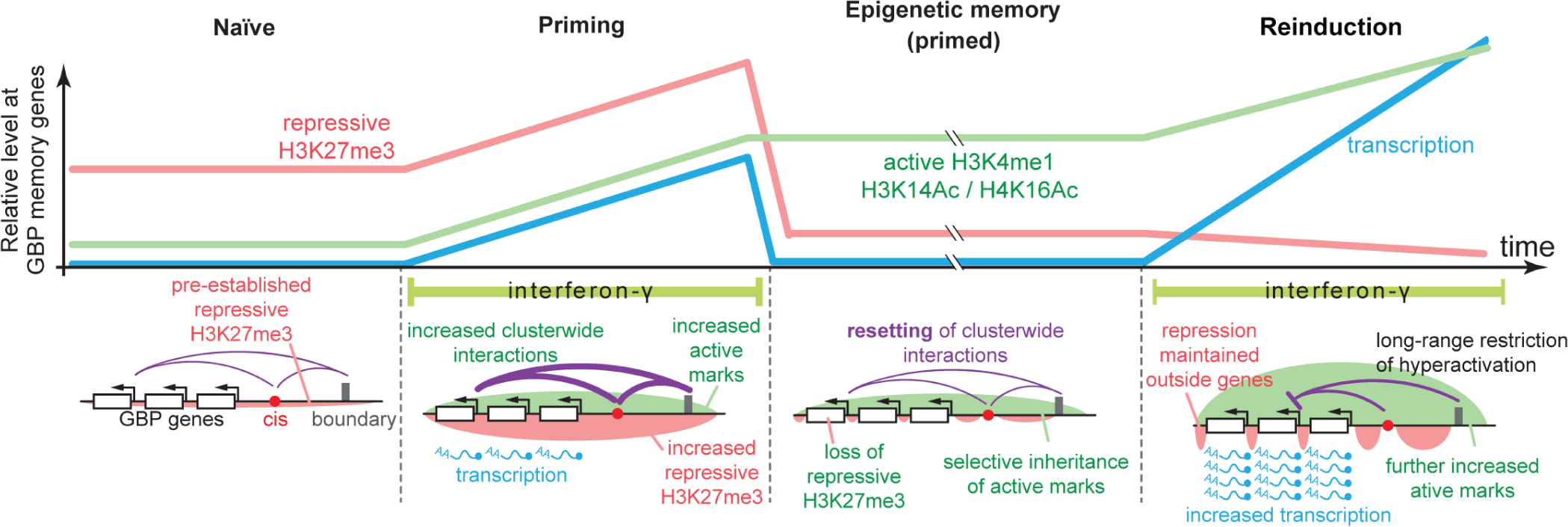
Proposed model for IFNγ-inducible chromatin-based transcriptional memory at GBP genes. The GBP cluster is embedded in a broad domain of low-level repressive H3K27me3 chromatin. IFNγ activation results in GBP transcription, and increased long-range interactions between the cis-regulatory elements, cluster boundaries and genes. It further results in establishing activating chromatin in part by KAT7, but also a further elevation of repressive chromatin mediated by PRC2. In the primed state, transcription is lost but active chromatin is selectively retained and mitotically heritable while suppressive H3K27me3 chromatin is locally depleted from GBP genes. This allows rapid and strong reactivation of GBP genes upon re-exposure to IFNγ. The cis-regulatory element acts to repress GBPs across the cluster preventing hyperactivation by IFNγ.

## Methods

### Cell culture

HeLa Kyoto cells (female, RRID: CVCL_1922) were grown in Dulbecco’s Modified Eagle Medium (DMEM) containing high glucose and pyruvate (ThermoFisher, 41966-029) supplemented with 10% NCS (newborn calf serum, ThermoFisher, 16010-159) and 1% Penicillin–Streptomycin (ThermoFisher, 15140-122) at 37°C, 5% CO2. For temporal depletion experiments using siRNAs or drugs, 1% Penicillin–Streptomycin has been omitted in DMEM. For passaging, cells were washed with 1× DPBS (ThermoFisher), detached with TrypLE Express phenol red (ThermoFisher), and resuspended in DMEM. Cells were counted using Countess™ Cell Counting according to the manufacturer’s instructions (Thermo Fisher Scientific). Transfection of cells was performed using Lipofectamine LTX (Thermo Fisher Scientific) according to the manufacturer’s instructions. Cells were routinely tested for Mycoplasma contamination.

### Transcriptional memory assay

Cells were primed with 50 ng/ml IFNγ (Merck) or left untreated for 24 h, followed by IFNγ washout with DPBS (ThermoFisher) and trypsinization by TrypLE (ThermoFisher) to harvest cells. Cells were cultured with fresh medium for another 48 h unless stated otherwise. Next, naïve and primed cells were induced by IFNγ for 24 h. After 24 h, cells were trypsinized and harvested, and the pellets were processed for subsequent experiments.

### Cut&Run-seq

Cut&Run-seq was performed using CUTANA v3 kit (Epicypher) according to the manufacturer’s protocol and with mild crosslinking (1 min incubation with 1% formaldehyde at room temperature). The antibodies used were: α-H3K4me1 (Epicypher, #13-0057), α-H3K4me3 (Epicypher, #13-0041), α-H3K14ac (Merck, #07-353), α-H4K16ac (Merck, #07-329), α-H3K27me3 (Cell Signaling Technology, #9733). Sequencing libraries were performed with NEBNext Ultra II DNA Library Prep Kit for Illumina (NEB) according to the published protocol^50^. The samples were multiplexed with NEBNext Multiplex Oligos for Illumina (Index Primers Set 1 and 2) (NEB). Size selection steps were performed with Ampure XP beads (Beckman Coulter) and adjusted for nucleosomal DNA fragment size (150bp, excluding adapters). The experiments were performed in biological duplicate or triplicate. The yield and quality of sequencing libraries were assessed by Qubit HS dsDNA Quantification Assay Kit (Thermo Fisher Scientific) and TapeStation 4150 System (Agilent). Multiplexed libraries were diluted to 1, 2 or 4 nM concentration and run on NextSeq 550 sequencer (Illumina) with NextSeq 500/550 High Output v2.5 (75 cycles PE) kit (Illumina).

### Expression (RT-qPCR)

Cell pellets (1 mln cells per sample) were re-suspended in 0.2 mL PBS and 0.8 mL TRIzol Reagent (Thermo Fisher Scientific). Cells were lysed by vortexing and incubated for 5 min at room temperature. Next, 0.16 mL chloroform was added per sample, mixed and incubated for 5 min at room temperature followed by centrifugation at 12000 g for 15 min at room temperature. The aqueous phase was mixed 1:1 (v:v) with 100% isopropanol and incubated at −20 C for 30 min, followed by centrifugation at 12000 g for 30 min at 4°C. The supernatant was removed and the pellet was washed with 1 mL of 75% ethanol and air-dried for 10 minutes. Finally, RNA pellets were re-suspended in 50 μL nuclease-free water. Any residual DNA contamination was removed with TURBO DNA-free™ Kit (Thermo Fisher Scientific), according to the manufacturer’s protocol. 1.5-2 ug RNA per sample was taken for cDNA synthesis, performed using a High-Capacity RNA-to-cDNA Kit (Applied Biosystems). Final cDNA samples were diluted 10 times before qPCR measurements. The qPCR assay was performed with iTaq Universal SYBR Green Supermix (Biorad), according to the manufacturer’s protocol. All RT-qPCR assays were performed in technical and biological triplicates. Primers used are specified in Table S1. Primer efficiency was determined computationally from amplification efficiency per PCR cycle using LinReg software^51^. The qPCR conditions were: 95°C 3 minutes; [95°C 10 s; 60°C 30 s]x50 cycles, followed by melting curve step (temperature range: 95-60C). The relative expression level of target genes was calculated using the efficiency-corrected ΔΔCt method (Pfaffl method)^52^.

### RNA interference and small molecule inhibitors siRNA

All siRNA transfections were performed on trypsinized cells. The cells were seeded in 6-well plates (2.25 x 10^5 cells per well) supplemented with 5 nM siRNA premixed with Opti-MEM Reduced Serum Medium (Gibco) and Lipofectamine RNAi Max Transfection Reagent (Thermo Fisher Scientific), according to the manufacturer’s protocol. The siRNAs were obtained from Silencer Select Pre-Designed and Validated siRNA (Thermo Fisher Scientific): KAT7 siRNA-1 (108177), KAT7 siRNA-2 (108179). Neg9 (N9) depletion siRNA target 5′-UACGACCGGUCUAUCGUAGTT -3′ was used as a control. The experiments were performed in biological triplicate.

### Small molecule inhibitors (EZHi and DOT1Li)

EZHi (UNC1999) and DOT1Li (SGC0946) were obtained from the Structural Genomics Consortium (SGC)^24^. The incubation with inhibitors was performed on trypsinized cells. GBP1-GFP cells were seeded in 24-well plates (1.6 x 10^4 cells per well) in 1 ml Dulbecco’s Modified Eagle Medium (DMEM) supplemented with 2 μM of the respective inhibitor. Mock control (cells supplemented with 100% DMSO in the same volume as inhibitors) was included in each experiment. The experiments were performed in 3 biological replicates per condition (priming, reinduction, naïve cells). For harvesting, cells were washed with PBS, trypsinized, crosslinked (1% formaldehyde, 10 min on rotator at room temperature) and quenched with 0.25 M glycine. The cells were subjected to FACS according to the description below. The experiments were performed in biological triplicate.

### GBP1-GFP line generation

The GBP1-GFP HeLa cell line was constructed using the LentiCRISPR V2-Blast (Addgene #83480) vector containing Cas9 sequence and a gRNA targeting exon 11 of the GBP1 gene that encodes the stop codon (See Table S1 for gRNA sequence). The homology repair template cloned in pUC19 consisted of a synthesised P2A-GFP cassette (Life Technologies) flanked by the GBP1 homology arms that match coordinates: chr1: 89053857-89053357 and 89053357-89052857. A silent mutation in the protospacer-adjacent motif (PAM) recognition sequence was introduced in the gRNA target of the homology arm to prevent Cas9 re-cutting after successful repair. The homology repair template was linearised before reverse co-transfection with the plasmid containing the Cas9/gRNA (1:3 ratio) using LipofectamineTM 3000 (Thermo Fisher Scientific). The following day, cells were subjected to blasticidin treatment for 48 h. Subsequently, cells were induced with IFNγ for 24 h and sorted to single cells by FACS based on the GFP fluorescence to generate monoclonal lines. Cells were maintained in culture for at least two weeks to erase IFNγ priming before being used in experiments.

### High-content microscopy screening of small molecules

Small molecule inhibitors were obtained from the Structural Genomics Consortium (SGC) as SGC Epigenetic Chemical Probes library^24^. Incubation with inhibitors was performed on trypsinized cells. GBP1-GFP cells were seeded in 96-well plates (1.6 x 10^4 cells per well) in 0.2 ml Dulbecco’s Modified Eagle Medium (DMEM) supplemented with 2 μM of the respective inhibitor. Mock control (cells supplemented with 100% DMSO in the same volume as inhibitors) was included in each experiment. The experiments were performed in 3 biological replicates per condition (priming, reinduction, naïve cells). Border wells of the plate were filled with PBS and excluded from the experiment to prevent temperature effects on the readout. Following transfection, the plates were incubated at room temperature for 45 min followed by standard growth conditions. For harvesting, cells were washed with PBS, crosslinked (1% formaldehyde, 10 min on rotator at room temperature) and quenched with 0.25 M glycine. Next, the cells were washed with PBS and stored at 4 C for up to 7 days before imaging as microscopy-high throughput screening (microscopy-HTS). Microscopy-HTS was performed on an Opera Phenix Plus High-Content Screening System (Perkin Elmer). GFP fluorescence thresholds were adjusted per plate based on the naïve (non-fluorescent) and priming (fluorescent) conditions. Final threshold per plate was selected based on the Z-score between conditions. The percentage of cells above the threshold was used to compare controls and inhibitor-treated samples to select hits affecting GBP1-GFP expression.

### Live-cell imaging

GBP1-GFP reporter cells (described above) were transduced with a pBABE retrovirus expressing H2B-mRFP^53^ to mark nuclei, facilitating analysis. Clones were selected by puromycin resistance and scored for robust H2B-mRFP expression. Cells were primed as per “Transcriptional memory assay” described above. 5 days after IFNy washout (memory window) cells were transferred into the chambers of a µ-Dish 35 mm Quad dish (Ibidi) with polymer coverslip and cultured for 24 hours in CO_2_-independent Live Cell Imaging Solution (Invitrogen) supplemented with 10% FBS (Life Technologies). Cells were then either induced with 50 ng/ml IFNγ or left untreated and imaged at 1-hour intervals for 24 hours, commencing 40-60 minutes after the addition of IFNy. Cells were imaged on a temperature-controlled Leica DMI6000 widefield microscope at 37°C equipped with a Hamamatsu Flash Orca 4.0 sCMOS camera, using a 40× 1.4 NA objective (HC PLAN APO). GFP fluorescence was quantified based on nuclei detection using TrackMate Cellpose plugin for ImageJ. The time-lapse tracks were analysed in R and filtered for continuous tracks to exclude those shorter than 24 hours. Each track was then normalised based on the first three time points. Datapoints of cells transitioning through mitosis were excluded due to transient increase in background fluorescence. The resulting tracks were used to determine the cut-off value for cells with GBP1-GFP expression or hyperactivated expression. To create a stringent cut-off, cells were considered expressing if GFP fluorescence was ≥ 3 times the interquartile range above the third quartile of the GFP signal in naïve cells. Similarly, cells were considered as hyperactivated expression if GFP fluorescence was ≥ 3 times the interquartile range above the third quartile of the GFP signal in cells during priming.

### FACS

For fluorescence-activated cell sorting (FACS) and cytometry, cells were collected by centrifugation for 5 min at 500 g, re-suspended in ice-cold Sorting Medium (1% Fetal Bovine Serum in PBS, 0.25mg/mL Fungizone (Thermo Fisher Scientific), 0.25μg∕mL/10μg∕mL Amphotericin B/Gentamicin (GIBCO)) and filtered using 5 mL polystyrene round-bottom tubes with cell-strainer caps (Falcon) before sorting and cytometry on FACSAria III Cell Sorter (BD Biosciences). For sorting, the cells were collected in 96-well plates with Conditional Medium (1:1 mixture of fresh complete medium and medium collected from proliferating cell cultures that is 0.45μm filtered, supplemented with 20% Fetal Bovine Serum, 0.25mg/mL Fungizone (Thermo Fisher Scientific), 0.25μg∕mL/10μg∕mL Amphotericin B/Gentamicin (GIBCO)).

### CRISPR/Cas9 cloning and genome engineering

E1, E2-1 and E2-2 mutants were generated with CRISPR/Cas9 technology as double-cut cis-regulatory elements’ deletion lines. The gRNAs were designed using IDT and CRISPick (Broad Institute) tools. The gRNA sequences are specified in Table S1. Relevant gRNA pairs were cloned into lentiCRISPR v2-Blast (Addgene #83480) and lentiCRISPR v2 (Addgene #52961) to allow for dual antibiotic resistance after transfection (blasticidin and puromycin, respectively).

Resultant plasmids were co-transfected with viral packing plasmid psPAX2 (Addgene #12260) and viral envelope plasmid pMD2.G (Addgene #12259) into HEK293T cells at a molar ratio of 4:3:1, respectively followed by incubation at 37C for 3 days. Culture medium containing lentiviral particles was collected, filtered through 0.45µm filters, incubated with 8 mg/mL Polybrene Reagent (Merck) for 1h, mixed 1:1 with fresh medium and added to Hela cells for transduction. 2 days after transduction, the cells drug selected with 5 mg/mL blasticidin and 1 mg/ml puromycin. Mutant lines were collected and validated with gDNA PCR and Sanger sequencing using the oligonucleotides specified in Table S1. E2-2 monoclonal line was generated by single-cell FACS as described above.

### Genome architecture (Capture-C-seq and 3C-qPCR)

Capture-C-seq was performed as published^34^ with the following specifications. The viewpoints were selected and their specific probes were designed using Capsequm2 software. 5 x 10^6 cells were used per sample. DNA was digested with DpnII restriction enzyme (NEB) and religated with T4 DNA HC ligase (Thermo Fisher Scientific). Sonication was performed on Q500 machine (QSonica) to obtain ∼200bp DNA fragments with pre-optimization of sonication conditions performed on genomic DNA control samples. Sequencing libraries were synthesized and multiplexed with NEBNext Ultra II kit (NEB). Ampure XP (Beckman Coulter) were used for size selection according to the manufacturer’s protocol. The experiment was adapted for high-specificity sequencing (double hybridization with probe titration). The hybridization was performed in two separate pools with 5’-biotinylated oligonucleotides for either, E2 or CH-C, viewpoint. The oligonucleotides are listed in Table S1. The experiments per each pool were performed in biological duplicate. Sequencing was performed on NextSeq550 sequencer (Illumina) using NextSeq 500/550 High Output Kit v2.5 (75 Cycles PE) (Illumina).

3C-qPCR was performed according to the published protocol^54^ with the following specifications. 0.8-1 x 10^6 cells were used per sample. DNA digestion was performed with DpnII restriction enzyme (NEB) and religated with T4 DNA HC ligase (Thermo Fisher Scientific). Removal of residual proteins and RNA was performed by Proteinase K (Ambion) and RNase A (Thermo Fisher Scientific) treatments, according to the manufacturer’s protocols. DNA was purified with phenol-chloroform-isoamyl alcohol and ethanol, according to the published protocol^55^. Sample yield and quality were assessed by gel electrophoresis, Qubit BR dsDNA assays (Thermo Fisher Scientific) and Real-Time quantitative PCR (RT-qPCR) analyses performed on genomic, digested and re-ligated controls. For RT-qPCR assays, each final 3C sample was diluted to 25 ng DNA per reaction. Ct values were normalized according to the published protocol^54^ with primer efficiency determined computationally from amplification efficiency per PCR cycle using LinReg software^51^ and amplification of E2 or CH-C baits within digested fragment set as a loading control. The oligonucleotides used for RT-qPCR are listed in Table S1. The experiments were performed in biological triplicate.

### Bioinformatic data analysis and statistics Unix

All unix commands were performed in conda environments. For Cut&Run-seq data analysis, raw reads (fastq) per experiment were downloaded from Basespace servers (Illumina) using basespace-cli and concatenated per sample using base unix. Read quality was assessed using Basespace (Illumina) and FastQC^56^ software. Next, reads were mapped to hg38 genome with Bowtie2^57^, adjusting trimming conditions dependent on the read quality. SAM to BAM conversion, BAM sorting and indexing were performed with Samtools v1.1^58^. Read duplicates were removed with Picard (MarkDuplicates command)^59^. Sorted and duplicate-removed BAM files were subjected to read count, normalization (CPM) and conversion to bigwig format with Deeptools v2 (bamcoverage command)^60^. Bigwig files were visualized in IGV^61^ and WashU Epigenome Browser^62^. The read count matrices for cross-comparison between conditions and samples were generated with Deeptools v2 (multibigwigsummary command)^60^.

For Capture-C-seq analysis, raw reads (fastq) per experiment were downloaded from Basespace servers (Illumina) using basespace-cli and concatenated per sample using base bash. Read quality was assessed using Basespace (Illumina) and FastQC^56^ software. Next, reads were processed with CapCruncher pipeline^63^ up to the generation of compressed contact matrices (HDF5 format). Contact matrices were further processed and converted to bedpe format with Cooler tool^64^. Filtering and normalization was performed with custom-made scripts in base unix using the same method as in the published protocol^63^. Final bedpe or bedgraph files were visualized in IGV^61^ and WashU Epigenome Browser^62^. The read count matrices for cross-comparison between conditions and samples were generated with Deeptools v2 (multibigwigsummary command)^60^.

### R

The read count matrices (Deeptools v2 multibigwigsummary ouput) from Cut&Run-seq and Capture-C-seq were processed in R v4.1^65^ with RStudio v1.4^66^ interface. The matrices were annotated, filtered (removal of 0 count reads) and quantified (mean, standard error, folds between conditions and statistics) using custom-made scripts. Data wrangling was performed using base R and dplyr package^67^. Data visualization was performed using ggplot2^68^ and ggrepel^69^ packages.

### Statistics

If not specified otherwise, the statistics for pairwise comparison between conditions or samples were performed using Student’s T-test in Microsoft Excel or base R. Hetero- or homoscedasticity was determined using F-test in Microsoft Excel or base R. The relevant significance levels are plotted as tabulated *P* values in each figure The error bars on bar plots throughout the paper correspond to SEM.

## Data availability

All sequencing data, raw reads and processed files, were deposited in Gene Expression Omnibus (GEO) and will be publicly accessible after peer review.

## Code availability

The scripts for the bioinformatic analyses with their parameters were deposited in public repository on GitHub under the link: https://github.com/Pwmski/mikulski-lab/tree/main/Mikulski-et-al-2023

## Acknowledgements

We thank members of the Jansen lab and Wojciech Siwek (University of Gdańsk) for helpful discussions. We thank Jim Huges and Damien Downes (University of Oxford) for advice on Capture-C. We thank the Structural Genomics Consortium for access to the Epigenetic Chemical Probes library and members of Klose, Barr, Brockdorff, Nasmyth and Gergely labs (University of Oxford) for support and fruitful exchanges. This work was funded by a Senior Wellcome Research Fellowship 210645/Z/18/Z to LETJ and a Goodger and Schorstein Scholarship from the University of Oxford to PM. SSHT was supported by Fundação para a Ciência e a Tecnologia (FCT) doctoral fellowship PD/BD/128438/2017. AK is a Clarendon Scholar supported by the Hill Foundation.

## References

1. Margueron, R. & Reinberg, D. Chromatin structure and the inheritance of epigenetic information. Nat. Rev. Genet. 11, 285–296 (2010).

2. D’Urso, A. & Brickner, J. H. Epigenetic Transcriptional Memory. Curr. Genet. 63, 435 (2017).

3. Tehrani, S. S. H., Kogan, A., Mikulski, P. & Jansen, L. E. T. Remembering foods and foes: emerging principles of transcriptional memory. Cell Death and Differentiation (2023) doi:10.1038/s41418-023-01200-6.

4. Ivashkiv, L. B. IFNγ: signalling, epigenetics and roles in immunity, metabolism, disease and cancer immunotherapy. Nat. Rev. Immunol. 2018 189 18, 545–558 (2018).

5. Kamada, R. et al. Interferon stimulation creates chromatin marks and establishes transcriptional memory. Proc. Natl. Acad. Sci. U. S. A. 115, E9162–E9171 (2018).

6. Light, W. H. et al. A Conserved Role for Human Nup98 in Altering Chromatin Structure and Promoting Epigenetic Transcriptional Memory. PLOS Biol. 11, e1001524 (2013).

7. Gialitakis, M., Arampatzi, P., Makatounakis, T. & Papamatheakis, J. Gamma Interferon-Dependent Transcriptional Memory via Relocalization of a Gene Locus to PML Nuclear Bodies. Mol. Cell. Biol. 30, 2046–2056 (2010).

8. Siwek, W., Tehrani, S. S. H., Mata, J. F. & Jansen, L. E. T. Activation of Clustered IFNγ Target Genes Drives Cohesin-Controlled Transcriptional Memory. Mol. Cell 80, 396–409.e6 (2020).

9. Tretina, K., Park, E. S., Maminska, A. & MacMicking, J. D. Interferon-induced guanylate-binding proteins: Guardians of host defense in health and disease. J. Exp. Med. 216, 482–500 (2019).

10. Tehrani, S. S. et al. STAT1 is required to establish but not maintain interferon-γ-induced transcriptional memory. EMBO J. 42, e112259 (2023).

11. Wang, Z. et al. Combinatorial patterns of histone acetylations and methylations in the human genome. Nat. Genet. 2008 407 40, 897–903 (2008).

12. Regadas, I. et al. A unique histone 3 lysine 14 chromatin signature underlies tissue-specific gene regulation. Mol. Cell 81, 1766–1780.e10 (2021).

13. Taylor, G. C. A., Eskeland, R., Hekimoglu-Balkan, B., Pradeepa, M. M. & Bickmore, W. A. H4K16 acetylation marks active genes and enhancers of embryonic stem cells, but does not alter chromatin compaction. Genome Res. 23, 2053–2065 (2013).

14. Shogren-Knaak, M. et al. Histone H4-K16 acetylation controls chromatin structure and protein interactions. Science (80-.). 311, 844–847 (2006).

15. Skene, P. J. & Henikoff, S. An efficient targeted nuclease strategy for high-resolution mapping of DNA binding sites. Elife 6, (2017).

16. Blackledge, N. P. & Klose, R. J. The molecular principles of gene regulation by Polycomb repressive complexes. Nat. Rev. Mol. Cell Biol. 2021 2212 22, 815–833 (2021).

17. Wutz, G. et al. ESCO1 and CTCF enable formation of long chromatin loops by protecting cohesinstag1 from WAPL. Elife 9, (2020).

18. Rao, S. S. P. et al. A 3D map of the human genome at kilobase resolution reveals principles of chromatin looping. Cell 159, 1665–1680 (2014).

19. Kueh, A. J. et al. Stem cell plasticity, acetylation of H3K14, and de novo gene activation rely on KAT7. CellReports 42, 111980 (2023).

20. MacPherson, L. et al. HBO1 is required for the maintenance of leukaemia stem cells. Nature 577, 266–270 (2020).

21. Ezhkova, E. et al. EZH1 and EZH2 cogovern histone H3K27 trimethylation and are essential for hair follicle homeostasis and wound repair. Genes Dev. 25, 485 (2011).

22. Shen, X. et al. EZH1 mediates methylation on histone H3 lysine 27 and complements EZH2 in maintaining stem cell identity and executing pluripotency. Mol. Cell 32, 491 (2008).

23. Xu, B. et al. Selective inhibition of EZH2 and EZH1 enzymatic activity by a small molecule suppresses MLL-rearranged leukemia. Blood 125, 346 (2015).

24. Brown, P. J. & Müller, S. Open access chemical probes for epigenetic targets. Future Med. Chem. 7, 1901 (2015).

25. Nguyen, A. T. & Zhang, Y. The diverse functions of Dot1 and H3K79 methylation. Genes Dev. 25, 1345–1358 (2011).

26. Lin, C. et al. AFF4, a Component of the ELL/P-TEFb Elongation Complex and a Shared Subunit of MLL Chimeras, Can Link Transcription Elongation to Leukemia. Mol. Cell 37, 429–437 (2010).

27. Guccione, E. et al. Methylation of histone H3R2 by PRMT6 and H3K4 by an MLL complex are mutually exclusive. Nat. 2007 4497164 449, 933–937 (2007).

28. Ogryzko, V. V., Schiltz, R. L., Russanova, V., Howard, B. H. & Nakatani, Y. The Transcriptional Coactivators p300 and CBP Are Histone Acetyltransferases. Cell 87, 953–959 (1996).

29. Rice, J. C. et al. Histone Methyltransferases Direct Different Degrees of Methylation to Define Distinct Chromatin Domains. Mol. Cell 12, 1591–1598 (2003).

30. Fanucchi, S. et al. Immune genes are primed for robust transcription by proximal long noncoding RNAs located in nuclear compartments. Nat. Genet. 2018 511 51, 138–150 (2018).

31. Pascual-Garcia, P., Little, S. C. & Capelson, M. Nup98-dependent transcriptional memory is established independently of transcription. Elife 11, (2022).

32. Owen, J. A., Osmanović, D. & Mirny, L. Design principles of 3D epigenetic memory systems. Science (80-.). 382, (2023).

33. Tan-Wong, S. M., Wijayatilake, H. D. & Proudfoot, N. J. Gene loops function to maintain transcriptional memory through interaction with the nuclear pore complex. Genes Dev. 23, 2610–2624 (2009).

34. Downes, D. J. et al. Capture-C: a modular and flexible approach for high-resolution chromosome conformation capture. Nat. Protoc. 2022 172 17, 445–475 (2022).

35. Kim, T. K., Hemberg, M. & Gray, J. M. Enhancer RNAs: A Class of Long Noncoding RNAs Synthesized at Enhancers. Cold Spring Harb. Perspect. Biol. 7, a018622 (2015).

36. Margueron, R. et al. Role of the polycomb protein EED in the propagation of repressive histone marks. Nat. 2009 4617265 461, 762–767 (2009).

37. Hansen, K. H. et al. A model for transmission of the H3K27me3 epigenetic mark. Nat. Cell Biol. 2008 1011 10, 1291–1300 (2008).

38. Ragunathan, K., Jih, G. & Moazed, D. Epigenetic inheritance uncoupled from sequence-specific recruitment. Science (80-.). 348, (2015).

39. Bostick, M. et al. UHRF1 plays a role in maintaining DNA methylation in mammalian cells. Science (80-.). 317, 1760–1764 (2007).

40. Sharif, J. et al. The SRA protein Np95 mediates epigenetic inheritance by recruiting Dnmt1 to methylated DNA. Nat. 2007 4507171 450, 908–912 (2007).

41. Samata, M. et al. Intergenerationally Maintained Histone H4 Lysine 16 Acetylation Is Instructive for Future Gene Activation. Cell 182, 127–144.e23 (2020).

42. Liu, Y. et al. Widespread Mitotic Bookmarking by Histone Marks and Transcription Factors in Pluripotent Stem Cells. Cell Rep. 19, 1283–1293 (2017).

43. Zhao, R., Nakamura, T., Fu, Y., Lazar, Z. & Spector, D. L. Gene bookmarking accelerates the kinetics of post-mitotic transcriptional re-activation. Nat. Cell Biol. 13, 1295–1304 (2011).

44. Calo, E. & Wysocka, J. Modification of Enhancer Chromatin: What, How, and Why? Molecular Cell vol. 49 825–837 (2013).

45. Bleckwehl, T. et al. Enhancer-associated H3K4 methylation safeguards in vitro germline competence. Nat. Commun. 12, 1–19 (2021).

46. Cheng, Q. J. et al. NF-κB dynamics determine the stimulus specificity of epigenomic reprogramming in macrophages. Science (80-.). 372, 1349–1353 (2021).

47. Fisch, D. et al. PIM1 controls GBP1 activity to limit self-damage and to guard against pathogen infection. Science 382, eadg2253 (2023).

48. Borden, E. C. et al. Interferons at age 50: past, current and future impact on biomedicine. Nat. Rev. Drug Discov. 2007 612 6, 975–990 (2007).

49. Ochando, J., Mulder, W. J. M., Madsen, J. C., Netea, M. G. & Duivenvoorden, R. Trained immunity — basic concepts and contributions to immunopathology. Nat. Rev. Nephrol. 2022 191 19, 23–37 (2022).

50. Liu, N. Nan Liu 2021. Library Prep for CUT&RUN with NEBNext® Ultra^TM^ II DNA Library Prep Kit for Illumina® (E7645). protocols.io.

51. Ruijter, J. M. et al. Amplification efficiency: linking baseline and bias in the analysis of quantitative PCR data. Nucleic Acids Res. 37, e45 (2009).

52. Pfaffl, M. W. A new mathematical model for relative quantification in real-time RT–PCR. Nucleic Acids Res. 29, e45 (2001).

53. Bodor, D. L. et al. The quantitative architecture of centromeric chromatin. Elife 2014, (2014).

54. Rebouissou, C., Sallis, S. & Forné, T. Quantitative Chromosome Conformation Capture (3C-qPCR). Methods Mol. Biol. 2532, 3–13 (2022).

55. Green, M. R. & Sambrook, J. Precipitation of DNA with Isopropanol. Cold Spring Harb. Protoc. 2017, pdb.prot093385 (2017).

56. Babraham Bioinformatics - FastQC A Quality Control tool for High Throughput Sequence Data. https://www.bioinformatics.babraham.ac.uk/projects/fastqc/.

57. Langmead, B. & Salzberg, S. L. Fast gapped-read alignment with Bowtie 2. Nat. Methods 9, 357 (2012).

58. Danecek, P. et al. Twelve years of SAMtools and BCFtools. Gigascience 10, 1–4 (2021).

59. Institute, B. Picard Toolkit. GitHub Repos.

60. Ramírez, F. et al. deepTools2: a next generation web server for deep-sequencing data analysis. Nucleic Acids Res. 44, W160–W165 (2016).

61. Robinson, J. T. et al. Integrative Genomics Viewer. Nat. Biotechnol. 29, 24 (2011).

62. Li, D. et al. WashU Epigenome Browser update 2022. Nucleic Acids Res. 50, W774–W781 (2022).

63. Smith, A., Rue-Albrecht, K. & mtekman. sims-lab/CapCruncher: CapCruncher v0.3.8. (2023) doi:10.5281/zenodo.10116675.

64. Abdennur, N. & Mirny, L. A. Cooler: scalable storage for Hi-C data and other genomically labeled arrays. Bioinformatics 36, 311 (2020).

65. R Core Team. R: A Language and Environment for Statistical Computing. (2021).

66. RStudio Team. RStudio: Integrated Development Environment for R. (2020).

67. Wickham, H., François, R., Henry, L., Müller, K. & Vaughan, D. dplyr: A Grammar of Data Manipulation. (2023).

68. Wickham, H. ggplot2: Elegant Graphics for Data Analysis. (Springer-Verlag New York, 2016).

69. Slowikowski, K. ggrepel: Automatically Position Non-Overlapping Text Labels with ‘ggplot2’. (2023).

