## Supplementary figures and tables for "Transcriptional memory is conferred by combined heritable maintenance and local removal of selective chromatin modifications"

A

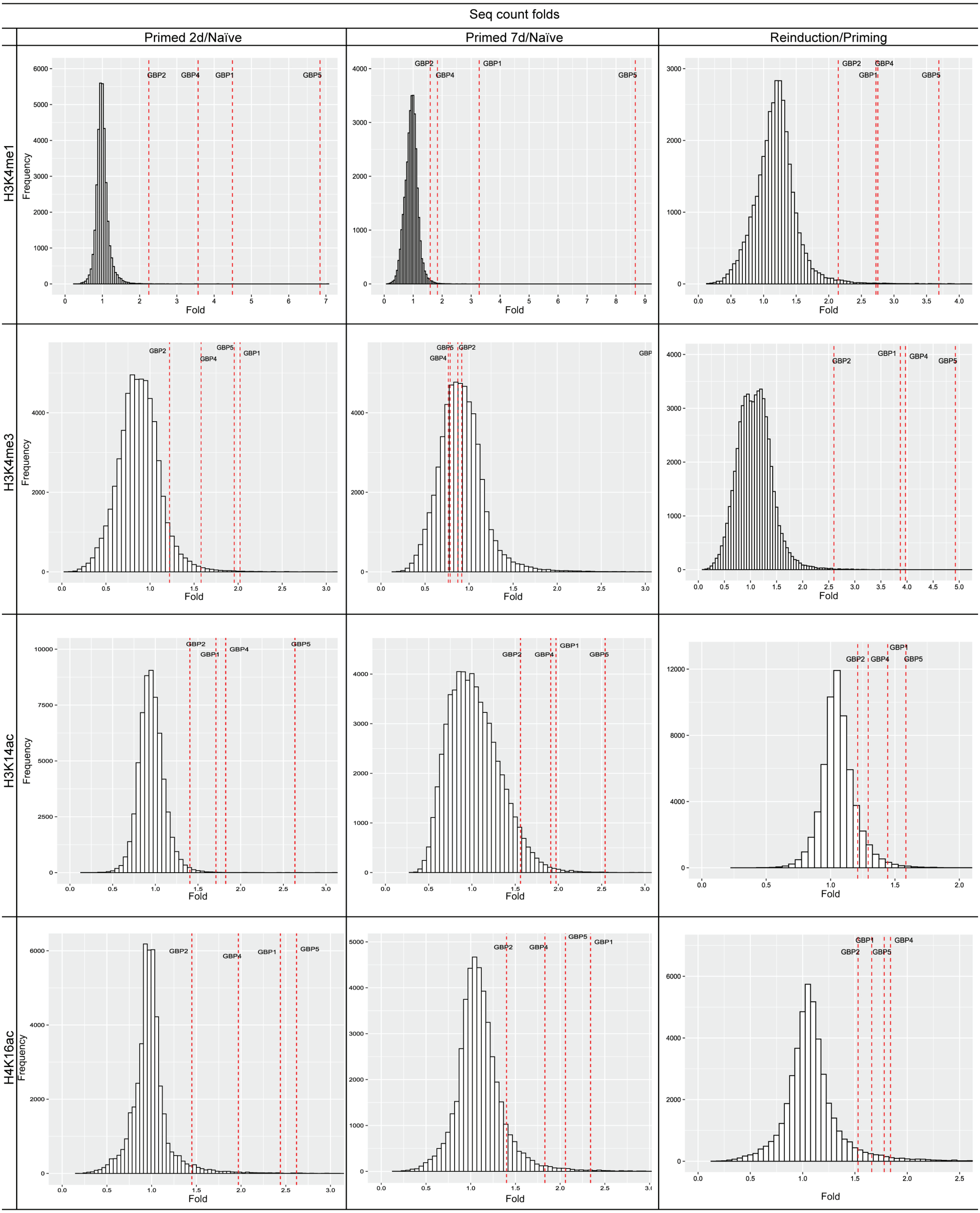


**Fig. S1. GBP memory genes exhibit selective retention of specific active chromatin modifications in IFNγ transcriptional memory on genome-wide scale**

**A.** Histograms for fold changes between indicated conditions for selected active chromatin modifications. The data corresponds to Cut&Run sequencing read count calculated genome-wide for each human gene (hg38 reference) with the enrichment over respective gene body and regions: 3kb upstream of transcriptional start site (promoter region) and 3kb downstream of transcriptional termination site (terminator region). Dashed vertical lines mark folds corresponding to GBP memory genes.

A

B

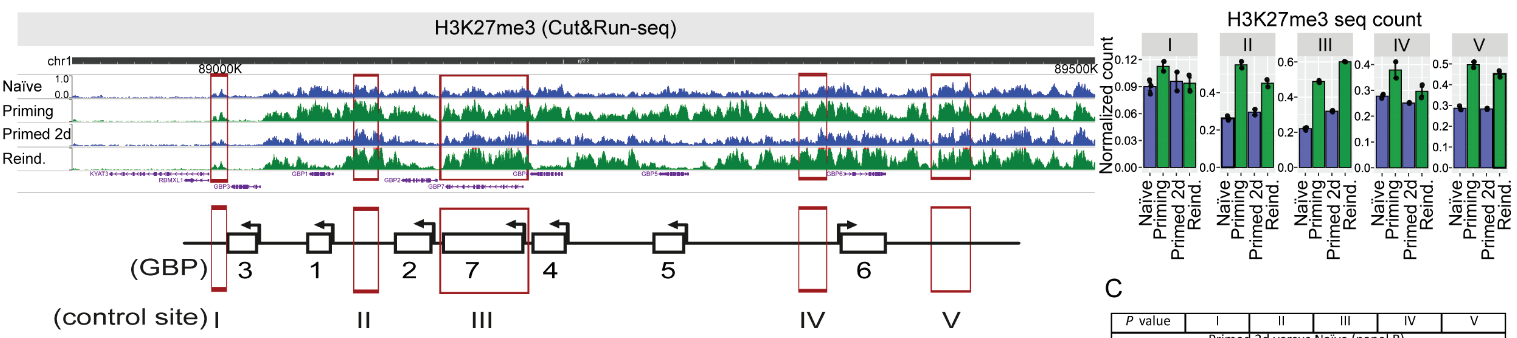


C

| *P* value | I | II | III | IV | V |
|---|---|---|---|---|---|
| Primed 2d versus Naïve (panel B) | | | | | |
| H3K27me3 | 3.0E-01 | 6.8E-01 | 4.8E-01 | 6.5E-02 | 3.4E-01 |
| Priming versus Naïve (panel B) | | | | | |
| H3K27me3 | 5.11E-02 | 3.71E-04 | 4.88E-05 | 2.46E-02 | 5.21E-04 |

D

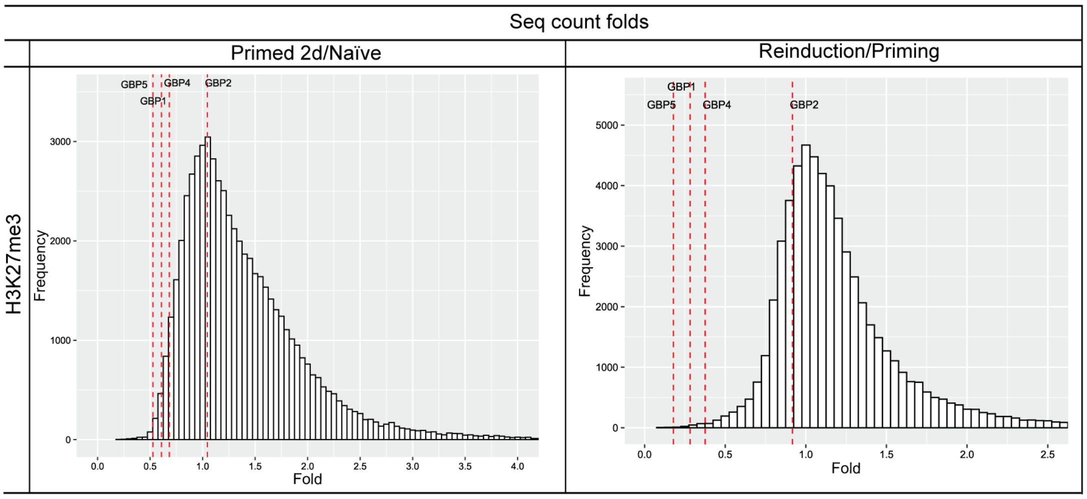


**Fig. S2. Repressive H3K27me3 chromatin is broadly accumulated across the GBP cluster but selectively removed from GBP genes in response to IFNγ**
**GBP cluster-wide analysis. A.** Cut&Run-seq enrichment of H3K27me3 in IFNγ-induced transcriptional memory regime represented as genome browser snapshots over GBP cluster. Red frames indicate regions outside of GBP memory genes broadly distributed in the cluster used for read quantification. **B.** Quantification of normalized Cut&Run sequencing reads for respective chromatin modifications over GBP cluster outside of GBP memory genes. The error bars correspond to SEM. The black dots on bar plots correspond to individual biological replicates. **C.** P values for selected pairwise comparisons for quantifications shown in panel B. P values ≤0.05 are highlighted in red. Statistical significance was calculated with two-sided t-test and prior determination of homo- or heteroscedasticity with F-test. **Gene-specific, genome-wide analysis. D.** Histograms of fold changes between indicated conditions for H3K27me3. The data corresponds to Cut&Run sequencing read count calculated genome-wide for each human gene (hg38 reference) with the enrichment over respective gene body and regions: 3kb upstream of transcriptional start site (promoter region) and 3kb downstream of transcriptional termination site (terminator region). Dashed vertical lines mark folds corresponding to GBP memory genes.

A

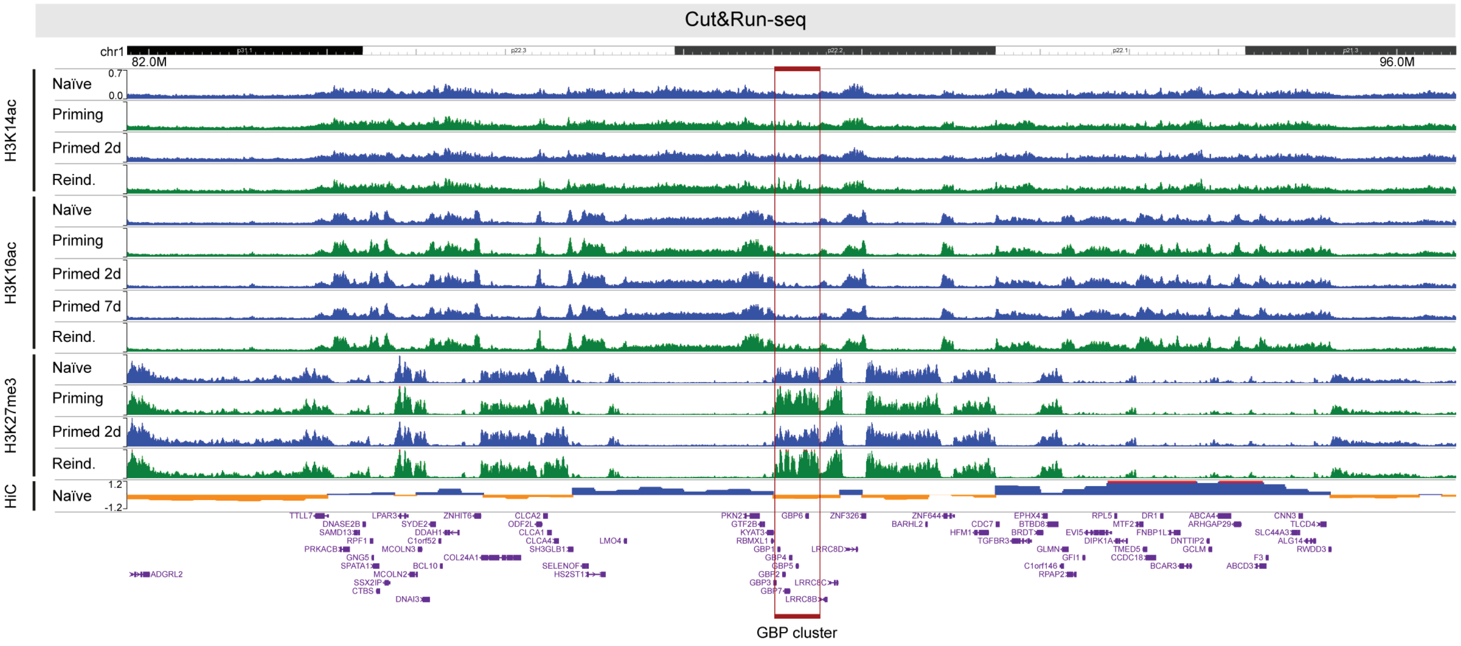


**Fig. S3. GBP cluster resides in a repressive topological chromatin domain**
**A.** Zoom-out chromosome visualization of Cut&Run-seq enrichment of selected active/repressive broad-distribution chromatin modifications and A/B compartmentalization (labelled blue and orange, respectively) from HiC assays (Wutz et al., EMBO J. 2017). Red frame indicates position of the GBP cluster.

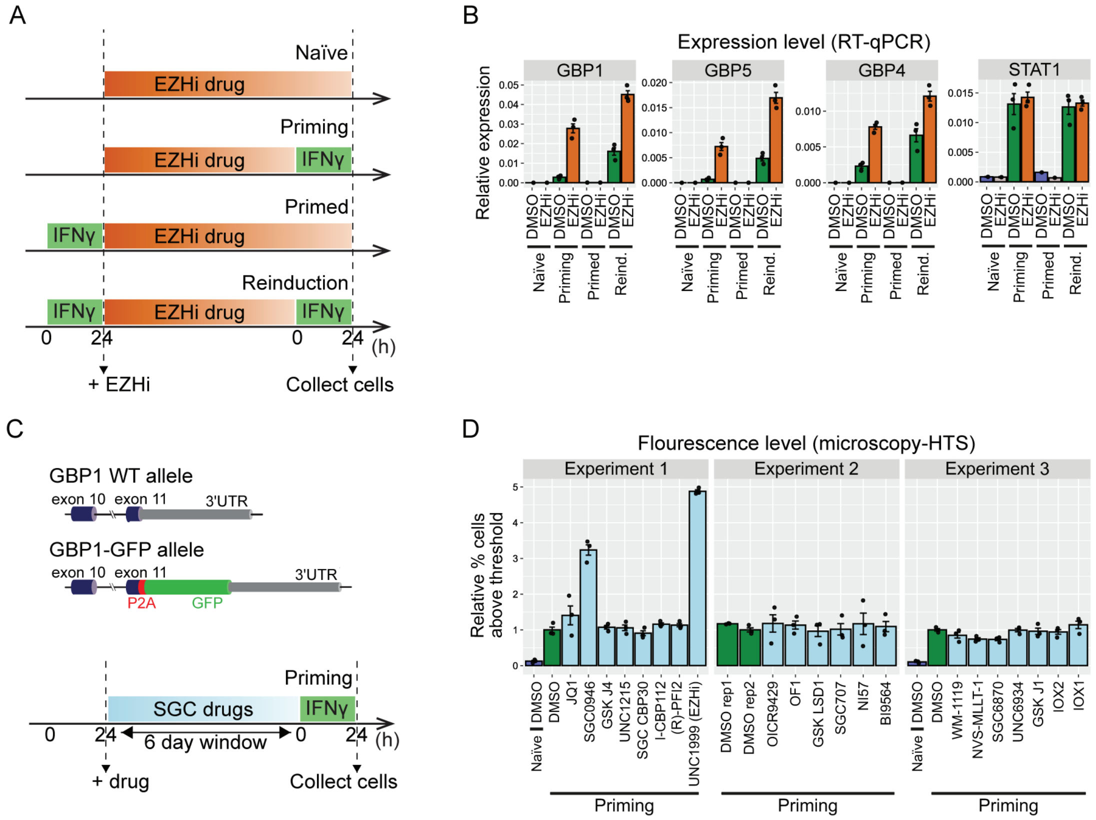


E

| *P* value | GBP1 | GBP5 | GBP4 | STAT1 |
|---|---|---|---|---|
| EZHi versus DMSO (panel B) | | | | |
| Priming | 9.2E-04 | 2.0E-03 | 5.2E-04 | 6.7E-01 |
| Reinduction | 9.9E-04 | 2.6E-02 | 1.8E-02 | 7.1E-01 |

| *P* value | Drug versus DMSO - Priming (panel D) | | | | | | | |
|---|---|---|---|---|---|---|---|---|
| | JQ1 | SGC0946 | GSK J4 | UNC1215 | SGC CBP30 | I-CBP112 | (R)-PFI | UNC1999 |
| Experiment 1 | 2.9E-01 | 3.9E-04 | 5.6E-01 | 6.9E-01 | 5.1E-01 | 2.1E-01 | 3.1E-01 | 4.0E-06 |
| | OICR 9429 | OF1 | GSK LSD1 | SGC707 | NI57 | BI9564 | | |
| Experiment 2 | 5.9E-01 | 4.3E-01 | 8.7E-01 | 9.4E-01 | 6.7E-01 | 6.5E-01 | | |
| | WM-1119 | NVS-MLLT-1 | SGC6870 | UNC6934 | GSK J1 | IOX2 | IOX1 | |
| Experiment 3 | 2.2E-01 | 1.4E-02 | 1.1E-02 | 9.4E-01 | 7.5E-01 | 5.4E-01 | 3.6E-01 | |

**Fig. S4. The expression and transcriptional memory of GBP genes is regulated by EZH1/2 and DOT1L**
**A.** Experimental scheme for temporal EZH1/2 depletion across IFNγ-stimulation regime. **B.** Expression changes of target genes upon temporal EZH1/2 depletion using EZHi drug compared to mock control (DMSO). The results correspond to delta-Ct values from RT-qPCR assays on cells harvested across IFNγ stimulation regime: naïve, priming, primed and reinduction conditions. **C.** Top: Schematic outline endogenous GBP1 gene structure in GBP1-GFP reporter line. Bottom: Experimental scheme for screening putative regulators for IFNγ transcriptional memory from SGC small molecule library. **D.** GBP1-GFP fluorescence intensity changes upon temporal drug-based depletion of screened regulators. The results correspond to % cells above fluorescence threshold normalized to DMSO control in priming condition. The threshold was adjusted per experiment based on fluorescence intensity in negative control – DMSO sample in non-IFNγ-induced, naïve cells. **E.** P values for selected pairwise comparisons for quantifications shown in panels: B and D. P values ≤0.05 are highlighted in red. Statistical significance was calculated with two-sided t-test and prior determination of homo- or heteroscedasticity with F-test. The error bars on all bar plots in the figure correspond to SEM. The black dots on bar plots correspond to individual biological replicates.

A

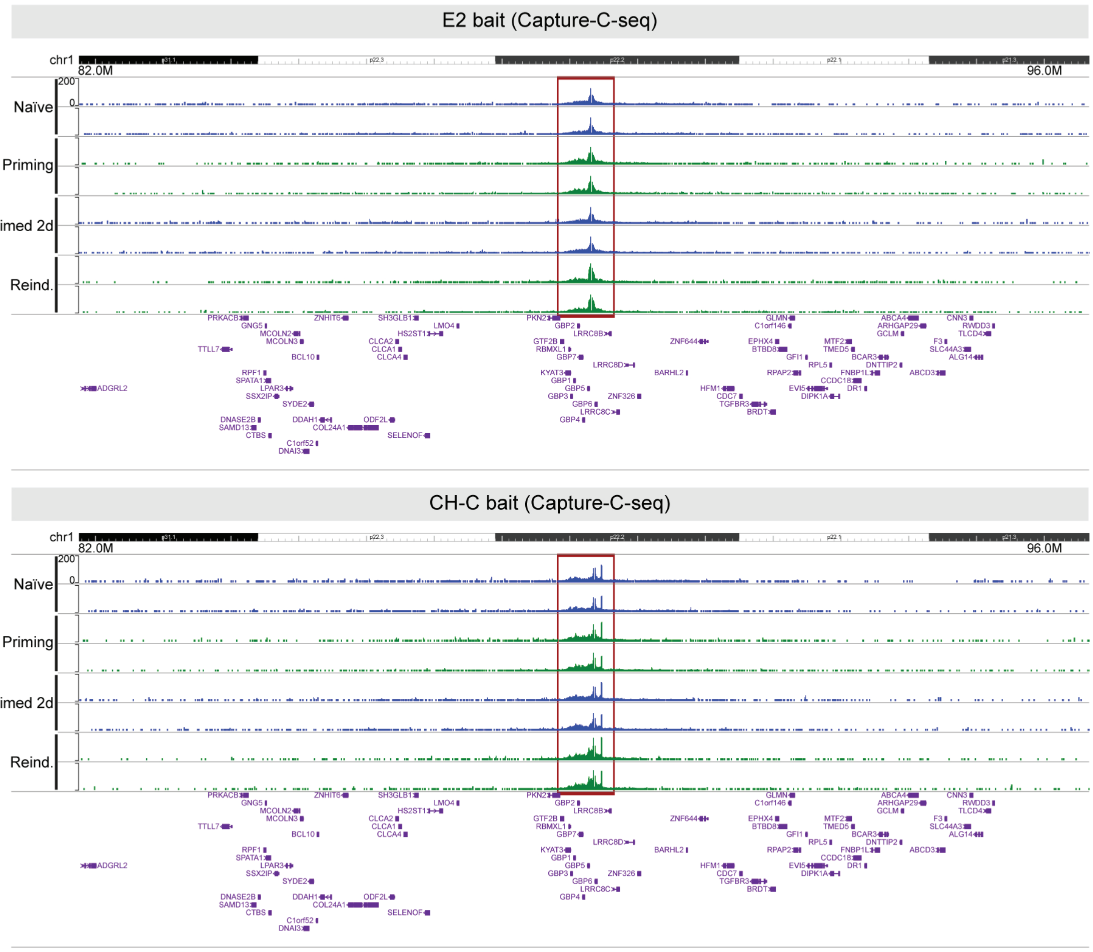


**Fig. S5. Long-range interactions from cis-regulatory elements E2 and CH-C are contained within GBP cluster only**

**A.** The enrichment of long-range interactions from element E2 (top panel) or Cohesin site CH-C (bottom panel) on zoom-out, near chromosome-wide scale. The results are presented as genome browser snapshots from normalized Capture-C sequencing reads at and around GBP cluster. Genome browser tracks show 2 biological replicates per condition in IFNγ-stimulation regime. Red frames indicate a position of GBP cluster.

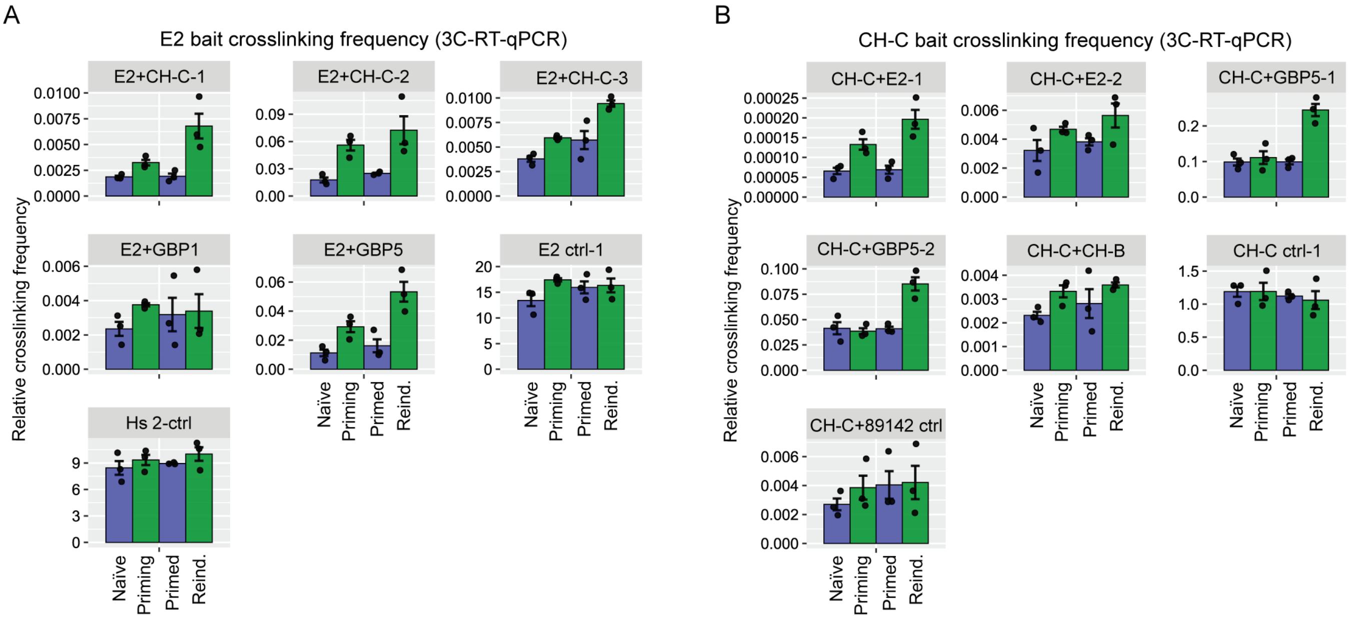


C

| *P* value | E2+CH-C-1 | E2+CH-C-2 | E2+CH-C-3 | E2+GBP1 | E2+GBP5 | E2 ctrl-1 | Hs 2-ctrl |
|---|---|---|---|---|---|---|---|
| Primed 2d vs Naïve (panel A) | | | | | | | |
| E2 bait | 8.7E-01 | 9.9E-02 | 1.8E-01 | 5.6E-01 | 4.7E-01 | 2.7E-01 | 6.2E-01 |

| *P* value | CH-C+E2-1 | CH-C+E2-2 | CH-C+GBP5-1 | CH-C+GBP5-2 | CH-C+CH-B | CH-C ctrl-1 | CH-C+89142 ctrl |
|---|---|---|---|---|---|---|---|
| Primed 2d vs Naïve (panel B) | | | | | | | |
| CH-C bait | 5.6E-01 | 8.4E-01 | 9.8E-01 | 9.5E-01 | 5.5E-01 | 5.4E-01 | 3.5E-01 |

**Fig. S6. IFNγ induces reciprocal long-range interactions between E2 and CH-C cis-regulatory elements**
**A.** The interaction level between E2 (bait) and target GBP cluster regions, including CH-C site. The results correspond to relative crosslinking frequency obtained from 3C-Real-Time quantitative PCR (RT-qPCR) assay. E2 and CH-C interaction is presented as 3 independent RT-qPCR amplicons (E2+CH-C 1/2/3). The results include control amplicons obtained with primers specific for the intact DNA fragment between DpnII sites after gDNA digestion – E2 bait (E2 ctrl-1) and Homo sapiens α-globin cluster (Hs 2-ctrl). **B.** The interaction level between CH-C (bait) and target GBP cluster regions, including E2 site. The results correspond to relative crosslinking frequency obtained from 3C-RT-qPCR assay. CH-C and E2 interaction is presented as 2 independent RT-qPCR amplicons (CH-C+E 1/2/3), complementing interactions shown in panel A. The results include control amplicons obtained with primers specific for the intact DNA fragment between DpnII sites after gDNA digestion – CH-C bait (CH-C ctrl-1) and Homo sapiens α-globin cluster (Hs 2-ctrl, presented in panel A). **C.** P values for selected pairwise comparisons for quantifications shown in panels: A and B. P values ≤0.05 are highlighted in red. Statistical significance was calculated with two-sided t-test and prior determination of homo- or heteroscedasticity with F-test. The error bars on all bar plots in the figure correspond to SEM. The black dots on bar plots correspond to individual biological replicates.

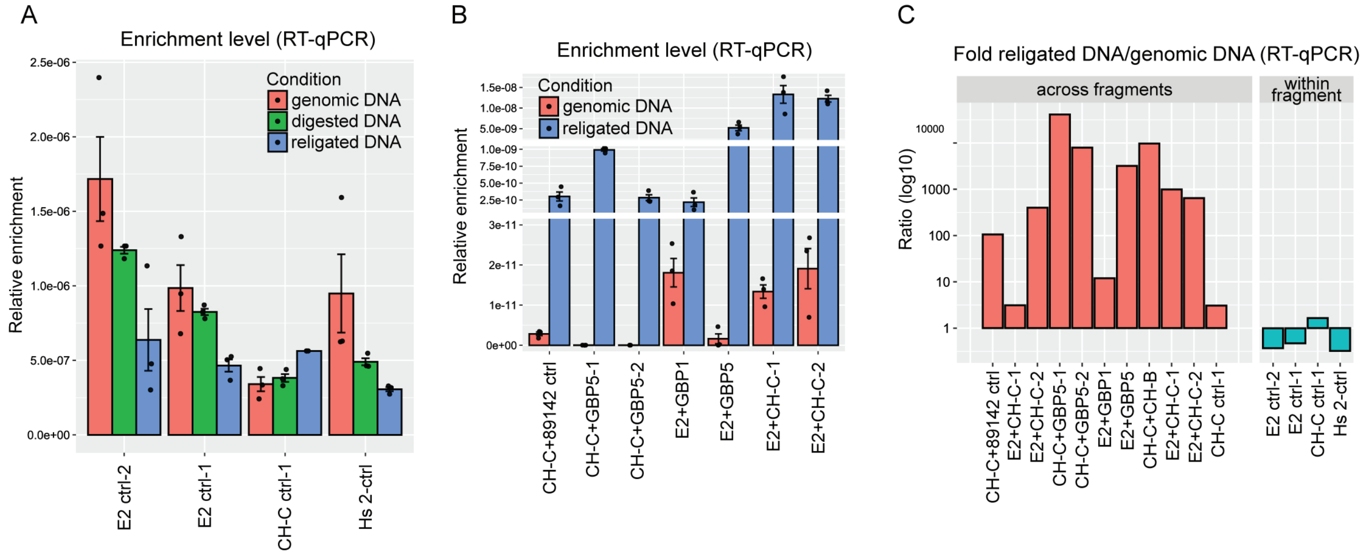


D

| *P* value | E2 ctrl-2 | E2 ctrl-1 | CH-C ctrl-1 | Hs 2-ctrl |
|---|---|---|---|---|
| Genomic DNA vs religated DNA (panel A) | | | | |
| Expression | 6.6E-02 | 5.6E-02 | 1.8E-01 | 6.3E-02 |

| *P* value | CH-C+89142 ctrl | CH-C+GBP5-1 | CH-C+GBP5-2 | E2+GBP1 | E2+GBP5 | E2+CH-C-1 | E2+CH-C-2 |
|---|---|---|---|---|---|---|---|
| Genomic DNA vs religated DNA (panel B) | | | | | | | |
| Expression | 2.1E-02 | 5.8E-04 | 3.2E-02 | 6.2E-02 | 3.4E-03 | 3.7E-02 | 6.4E-03 |

**Fig. S7. Technical validation of 3C-RT-qPCR assays.**
**A.** Enrichment level of controls specific for the intact DNA fragment between DpnII sites after gDNA digestion specified in Fig. S6. The results correspond to delta-Ct values from RT-qPCR assays in control samples before DNA digestion (genomic DNA), digested DNA and religated DNA. **B.** Enrichment level of selected interaction regions with E2 or CH-C baits specified in Fig. S6. The results correspond to delta-Ct values from RT-qPCR assays in control samples before DNA digestion (genomic DNA) and religated DNA. **C.** Assessment of religation efficiency and specificity of primers for target interactions. The results represent log(10)-transformed fold change between amplification of target regions in religated DNA to genomic DNA. The results correspond to log(10)-transformed delta-delta-Ct values from RT-qPCR assays in religated DNA and genomic DNA. The folds are presented for amplicons used in Fig. S6, grouped in categories: across DNA fragments after DpnII digestion (left group) or within given DNA fragment after DpnII digestion (right group). **D.** P values for selected pairwise comparisons for quantifications shown in panels: A and B. P values ≤0.05 are highlighted in red. Statistical significance was calculated with two-sided t-test and prior determination of homo- or heteroscedasticity with F-test. The error bars on all bar plots in the figure correspond to SEM. The black dots on bar plots correspond to individual biological replicates.

A

B

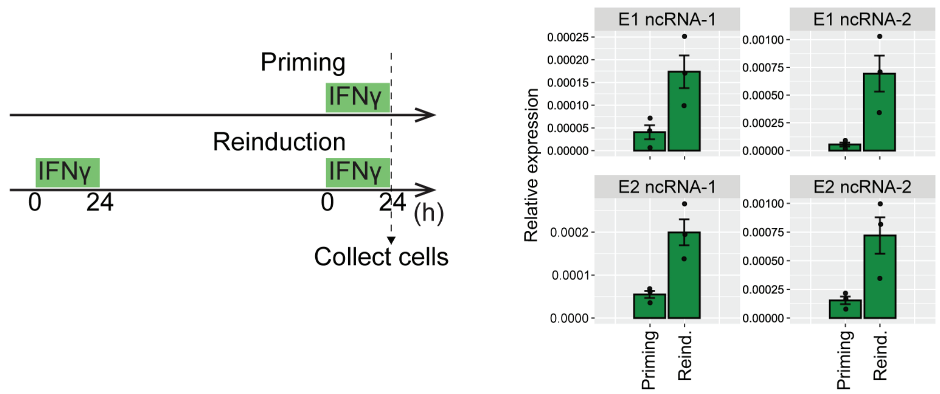


C

| *P* value | E1 ncRNA-1 | E1 ncRNA-2 | E2 ncRNA-1 | E2 ncRNA-2 |
|---|---|---|---|---|
| Priming vs Reinduction (panel B) | | | | |
| Expression | 5.0E-02 | 3.3E-02 | 2.0E-02 | 4.6E-02 |

D

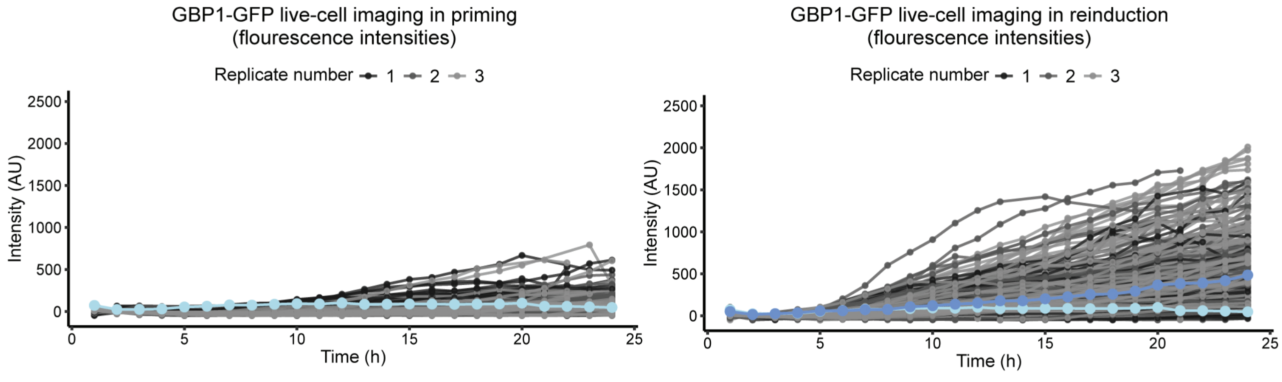


**Fig. S8 Cis-regulatory elements in GBP cluster produce ncRNAs showing transcriptional memory correlating with GBP expression dynamics**

**A.** Experimental transcriptional memory scheme and conditional cell harvesting regime. **B.** Expression levels of target cis-regulatory elements E1, E2 and CH-C in transcriptional memory timecourse. The results correspond to delta-Ct values from RT-qPCR assays on cells harvested in IFNγ priming and reinduction conditions. The results present expression levels of two independent (independent primer pairs) amplicons per cis-regulatory element. **C.** P values for selected pairwise comparisons for quantifications shown in panel B. P values ≤0.05 are highlighted in red. Statistical significance was calculated with two-sided t-test and prior determination of homo- or heteroscedasticity with F-test. The error bars on all bar plots in the figure correspond to SEM. The black dots on bar plots correspond to individual biological replicates. **D.** Time-lapse of live-cell GBP1-GFP protein expression during priming (left) and reinduction, 6 days after priming (right). Tracks of GFP intensities in individual cells were recorded over 24 hours of IFNγ exposure. Shades of grey represent cells from individual experiments. Light blue lines represent the threshold of GFP intensity set to quantify the fraction of cells with GBP1-GFP expression above baseline (naïve) levels. Dark blue line represent the threshold of GFP intensity set to quantify the fraction of cells with hyperactivated GBP1-GFP expression above the level observed in cells during priming.

| Table S1. Oligonucleotide sequences. | |
|---|---|
| **Name** | **Sequence 5'->3'** |
| **Expression (RT-qPCR)** | |
| GBP1 Fw | GTGGAACGTGTGAAAGCTGA |
| GBP1 Rv | CAACTGGACCCTGTCGTTCT |
| GBP5 Fw | TTCAATTTGCCCCGTCTGTG |
| GBP5 Rv | AGGCAGTGTTTCAAGTTGGG |
| GBP4 Fw | CAGTCTTCATGGAGCACTCCTTC |
| GBP4 Rv | TCAGGTGCTCTGAAAGCCGCTT |
| KAT7 Fw | GAATGCAAGGTGAGAGCACA |
| KAT7 Rv | CCGTGTGTTCCCATAGGTCT |
| STAT1 Fw | TGTATGCCATCCTCGAGAGC |
| STAT1 Rv | AGACATCCTGCCACCTTGTG |
| IRF1 Fw | TTTGTATCGGCCTGTGTGAATG |
| IRF1 Rv | AAGCATGGCTGGGACATCA |
| GBP2 Fw | AATTAGGGGCCCAGTTGGAAG |
| GBP2 Rv | AAGAGACGGTAACCTCCTGGT |
| CH-C ncRNA-1 Fw | GGTCTGGAGTCAAAGGAGAAAG |
| CH-C ncRNA-1 Rv | CCAGCTAGAAACAGTCCAACA |
| CH-C ncRNA-2 Fw | TCCCTGAATCTCTTCCTGTAAAC |
| CH-C ncRNA-2 Rv | TTGGGACAAGGCTGGATTT |
| E2 ncRNA-1 Fw | AGCTTTGAAGGGAACCAAGAG |
| E2 ncRNA-1 Rv | CTGGAAGCAAAGATAGTGAGTAGAG |
| E2 ncRNA-2 Fw | GTGTGCAATTTGGAGGGAATAAA |
| E2 ncRNA-2 Rv | AATGGCCAGCTGTTTGTAGA |
| E1 ncRNA-1 Fw | TAGACTTGAATTGGCCCAAGAG |
| E1 ncRNA-1 Rv | AGGAGGTGCTGGAAGTCTAA |
| E1 ncRNA-2 Fw | AGAGGAGGAGACTGCCAAA |
| E1 ncRNA-2 Rv | CCTATGCTTCTGTCTTCCTGTG |
| **Capture-C quality control (RT-qPCR)** | |
| CaptureC-cutsite Fw | GTCAGAAATAACAGGAAACCCAAA |
| CaptureC-cutsite Rv | TTACTTGTCGAACCCAGAAGAC |
| CaptureC-fragment Fw | GAGAATGGCCACATACAAGTAGA |
| CaptureC-fragment Rv | GGAGTTGTCAACACAAGCATATC |
| **Capture-C hybridization probes (5'-biotinylated)** | |
| E2 Left | GATCATTGGCCAGCTAATCTCAATTGGGCCAACACTTTTTCTCCTTTCTACACTTACTCATGGCACCCTGTATTAGTCCTTTACAATGCAGATAAGAAATTCAAATATTTCATTTGTTGG |
| E2 Right | ATGTTCTAACCAGAAAATATTGAAAAAGATTAATAAAAATAGATACACTTACCTACCCATTGATAAAGCAACATCAAACAAGAGACTGGGAAATTACCGTATGAGACAATGGATTAGATC |
| CH-C Left | GATCGGGAGATTCTCTTAATGAAAAGAAATTCAACAATTTTCCCTCTATCATGTGTGATTTTTTGGAGTCTTCTTAAAAACATCTTTCCCACACCAAGATAACAAAGATGCTCTCCTATA |
| CH-C Right | ACTGCCCACAGAGTCACTGGGGCAAAATGACCAACAAAACTGTAAAAGTGTGTCTTGTGGAAACAGCCATTTTAAAAAACGCTTTGATGGTGACAGGAATGCATGTGTCAACCGTGGATC |
| **3C-RT-qPCR E2 bait** | |
| E2 ctrl-1 Fw | GACTAGGCTCACAGACACATAAA |
| E2 ctrl-1 Rv | TCTGAGAGGTGAGAAACCTACTA |
| E2 ctrl-2 Fw | GAACCCTTGGCTGTCTCATT |
| E2 ctrl-2 Rv | CCTTGTCATCAGGAAGCAGTTA |

| | |
|---|---|
| E2+GBP1 Fw | AGAAACTGAGCACAGAAAGACT |
| E2+GBP1 Rv | TTGCTTAACTCTCTGTGGAGAAG |
| E2+GBP5 Fw | CACAGTGGTGACCAAATCAATAA |
| E2+GBP5 Rv | TGCTTAACTCTCTGTGGAGAAG |
| E2+CH-C-1 Fw | CTGATGACAAGGGATGCTGT |
| E2+CH-C-1 Rv | TCTGCTGCTGTTATCCCATTAG |
| E2+CH-C-2 Fw | CTTCCATCATTGTCTACCTCTGG |
| E2+CH-C-2 Rv | CCTTAGATATGCCCACATCCTG |
| Hs 2-ctrl Fw | GAGAATGGCCACATACAAGTAGA |
| Hs 2-ctrl Rv | GGAGTTGTCAACACAAGCATATC |
| E2+CH-C-3 Fw | AGACCAATCTTAACTGCTTCCT |
| E2+CH-C-3 Rv | AACTTTGCAATTGTGTCCTAGTC |
| **3C-RT-qPCR CH-C bait** | |
| CH-C ctrl-1 Fw | ACCATACAAGGAAATATTCAGAGGT |
| CH-C ctrl-1 Rv | TTCTCCATCATTGACCACTACAG |
| CH-C+E2-1 Fw | CAAAGACCAATCTTAACTGCTTCC |
| CH-C+E2-1 Rv | ACTGTGCTGCGTGTTAGG |
| CH-C+CH-B Fw | AGTCCTATCTTGCATCTATCATTCT |
| CH-C+CH-B Rv | AACTTTGCAATTGTGTCCTAGTC |
| CH-C+GBP5-2 Fw | CCTCCCTGCTTCCATTGTT |
| CH-C+GBP5-2 Rv | GGCTGAATGATGCTCCTTAGAT |
| CH-C+89142 ctrl Fw | AGGCCTGCCTACCCTTT |
| CH-C+89142 ctrl Rv | GAAACTTTGCAATTGTGTCCTAGTC |
| CH-C+GBP5-1 Fw | AAATCCTGTCTCCACACTATG |
| CH-C+GBP5-1 Rv | CCTCTGAATATTTCCTTGTATGGT |
| CH-C+E2-2 Fw | AGACCAATCTTAACTGCTTCCT |
| CH-C+E2-2 Rv | AACTTTGCAATTGTGTCCTAGTC |
| **GBP1-GFP cloning** | |
| GBP1 exon11 gRNA | CACCGACGAAAATGAGACGACGAA |
| **CRISPR E1/E2 deletion - PCR validation** | |
| E1 Fw | CACACTTCCTCGGGAGAACACTGAT |
| E1 Rv | ACAGTCTCTAACAGATCATCTCCCCG |
| E2 Fw | GGGGAGATAGGGCATACATGGGTAAAAG |
| E2 Rv | CTTCCTTCCTCACTCACCATGTACTCTG |
| **CRISPR E1/E2 deletion - sequencing validation** | |
| E2-2 Fw | CAACTATTCTGCTTTAGTACTTCG |
| E2-1 Fw | GGACTTCTTTACTATGCTAGATC |
| E2-2 Rv | CACATGGCTAGTCTCAAAATAGG |
| E1 Fw | AATTCCTGAGCTCCTATAACCCA |
| E1 Rv | CAATCGATCTCAGTAGAAGTCAG |
| **CRISPR E1/E2 deletion - gRNAs** | |
| E2-2 Left - oligo 1 | CACCGAAGACTAGACATTACCCAAT |
| E2-2 Left - oligo 2 | AAACATTGGGTAATGTCTAGTCTTC |
| E2-1 Left - oligo 1 | CACCGTGATCTCTCACCTGTTGTA |
| E2-1 Left - oligo 2 | AAACTACAACAGGTGAGAGATCAC |
| E2-1/2 Right - oligo 1 | CACCGAAATTACCGTATGAGACAA |
| E2-1/2 Right - oligo 2 | AAACTTGTCTCATACGGTAATTTC |
| E1 Left - oligo 1 | CACCGCCTAATTGCTGTACCAAGTT |
| E1 Left - oligo 2 | AAACAACTTGGTACAGCAATTAGGC |
| E1 Right - oligo 1 | CACCGCACGAATGTAGAGTCCAGCT |
| E1 Right - oligo 2 | AAACAGCTGGACTCTACATTCGTGC |
